# Outer membrane ‘PusCD’ complexes coordinate phospho(inositol)lipid utilisation in plant Bacteroidota

**DOI:** 10.64898/2026.09.03.749221

**Authors:** Alex N. Connolly, Laila Moushtaq, Pu Qian, Amanda A. Brindley, David J. Scanlan, Bert van den Berg, Andrew Hitchcock, Ian D.E.A. Lidbury

**Affiliations:** Molecular Microbiology: Biochemistry to Disease, School of Biosciences, University of Sheffield, Sheffield, UK; Plants, Photosynthesis and Soil, School of Biosciences, University of Sheffield, Sheffield, UK; School of Life Sciences, University of Warwick, Coventry, UK; Biosciences Institute, Faculty of Medical Sciences, Newcastle University, Newcastle upon Tyne, UK

## Abstract

Gram-negative bacteria belonging to the phylum Bacteroidota are prominent and ecologically important members of host-associated microbiomes. A feature of Bacteroidota species is the presence of diverse outer membrane TonB-dependent transporter and cognate surface-exposed lipoprotein (TBDT-SLP) complexes. Many ’SusCD-like‘ TBDT-SLPs coordinate complex carbohydrate capture from the environment. Here, we show the model soil Bacteroidota species *Flavobacterium johnsoniae* synthesises two TBDT-SLP complexes (named PusCD1 and PusCD2) that are essential for growth on phosphatidylinositol, a member of an abundant lipid class within plant plasma membranes, as a phosphorus source. Structural characterisation of ligand-bound PusCD1 complex from *F. johnsoniae* membranes reveals a new binding mechanism for organophosphorus and broadens the known functional repertoire of this class of transporters. Diverse PusCD-like complexes are widespread among plant-associated Bacteroidota, pointing to a major, previously unrecognised route for the recycling of organophosphorus compounds in the rhizosphere, with potential implications for engineering microbial-mediated phosphorus recovery from organic waste streams.

## Introduction

The outer membrane (OM) of Gram-negative bacteria is a semipermeable barrier permitting the diffusion of low molecular weight nutrients into the bacterial periplasm while maintaining membrane integrity and blocking the entry of larger compounds^1^. Consequently, the OM is incapable of establishing a proton motive force (PMF) to power the active import of larger or scarcely available nutrients. OM TonB-dependent transporters (TBDTs) are transmembrane beta-barrel proteins that address this problem through their association with the inner membrane (IM)-anchored energy transducing protein, TonB, utilising the PMF generated across the IM to drive substrate import at the cell surface^2–4^.

OM TBDTs are found in all Gram-negative bacteria known to date, being particularly abundant within the phylum Bacteroidota^4^. This radiation within the Bacteroidota may compensate for an inability to compete for scarce smaller nutrients in certain niches as members of this phylum often lack many IM ATP-binding cassette (ABC) or major facilitator superfamily-like transporters^5^. Many TBDTs in Bacteroidota are predicted to form a complex with a cognate surface-exposed lipoprotein (SLP) that contributes to ligand binding at the cell surface through a ‘pedal bin’ mechanism^6^. TBDT-SLP complexes are commonly encoded within gene clusters alongside carbohydrate-active enzymes (CAZymes) and surface glycan binding proteins, coordinating the binding, import, and breakdown of specific polysaccharide substrates^7^. These ‘polysaccharide utilisation loci (PUL)’ are a hallmark of Bacteroidota genomes, and are hypothesised to confer a fitness advantage in densely populated niches such as the human gut and the plant microbiome^8–10^ by capturing the diverse array of polysaccharides present in these environments^5,11–15^. The archetype PUL, named the ‘Starch utilisation system (Sus)’, gave rise to TBDT-SLP complexes being termed SusC (TBDT) and SusD (SLP)^16^. The vast diversity of SusCD-like pairs reflects their specificity towards individual glycans^4^, but may also indicate diversification of function beyond glycan import, playing roles in peptide^17,18^ and phosphorus (P)^7,19^ acquisition.

*Flavobacterium* and other genera belonging to Bacteroidota have emerged as important taxa within the plant microbiome^9,20^, and are significantly enriched in this niche compared to ‘bulk’ soil^10,21–23^. Most plant-associated *Flavobacterium* lack high-affinity ABC transporters for inorganic phosphate (P_i_), which is surprising given their ability to colonise plant roots, a that is niche commonly phosphate-limited due to plant uptake and high microbial activity^24,25^. We previously discovered that plant-associated *Flavobacterium* spp. synthesise SusCD-like pairs in response to phosphate depletion^19^. Unlike classical SusCD complexes encoded in PUL, these SusCD-like pairs were co-localised with phosphatases and P-responsive regulatory elements, forming discrete gene clusters that we termed ‘Phosphate utilisation systems (Pus)’. In *F. johnsoniae*, a total of four *pus* loci were identified of which two, *pus1* and *pus2*, encode putative ‘PusCD’ transport complexes that mimic canonical SusCD-like pairs^19^. *pus3* and *pus4* lack canonical SusCD-like pairs, instead containing TBDTs and alternative SLPs^19^. *pus1* encodes a putative phosphoinositide-specific phospholipase C (PI-PLC) and calcineurin-like phosphodiesterase, suggesting PusCD1 may specialise in phospholipid (PL)-derived organophosphate import.

Phospho(inositol)lipids constitute a major fraction of the plant plasma membrane (PM) and play crucial roles in the molecular cross-talk between the host and invading microbes^26–30^. The fate of root-derived PLs is unclear, though they are presumably in high abundance in the rhizosphere due to microbial cell turnover, root cell sloughing, and programmed cell death during root development^31–33^. Some plant-associated microbes are known to interfere with host phosphatidylinositol (PtdIn) signalling^34^, and PI-PLCs appear to be widespread across the genomes of terrestrial bacteria^35–37^. However, little is known as to whether these microbes use PLs as an alternative P source during growth under P-deplete conditions. In this study, we show that the PusCD transporter complexes encoded in *pus1* and *pus2* are essential for proper growth of plant-associated Bacteroidota on PtdIn and its corresponding headgroup glycerophosphoinositol (GroPI) as a sole P source. Using single-particle cryogenic electron microscopy (cryo-EM), we resolve a new mechanism for organophosphate binding and import at the cell surface.

## Results

### *Flavobacterium johnsoniae* grows on phosphatidylinositol as a phosphorus source

*F. johnsoniae* encodes several putative phospholipases and phosphatases required to hydrolyse the various ester bonds present in the PL headgroups of PtdIn, phosphatidylcholine (PtdChl) and phosphatidylethanolamine (PtdEth) (**Fig. 1A**). When grown in minimal medium, *F. johnsoniae* grew only on PtdIn as a sole P source and not on PtdChl or PtdEth (**Fig. 1B**) but grew on all three of the corresponding PL diester headgroups; GroPI, glycerophosphocholine (GroPC) and glycerophosphoethanolamine (GroPE) (**Fig. 1B**). The growth rate of the GroPI-grown cells was significantly greater (ANOVA: F = 203.3; d.f = 4,10; p <0.001, Tukey HSD p <0.001) than either GroPE- or GroPC-grown cells and was comparable to growth on P_i_ (**Fig. S1**). We next screened our previously characterised *F. johnsoniae* mutant (ΔM5) lacking four phosphomonoesterases, including PafA (Fjoh_0023) and PhoA1 (Fjoh_3249), which is unable to grow on P-monoesters as a sole P source^38^. ΔM5 failed to grow on GroPC and GroPE, but growth was restored by plasmid-borne complementation with either *pafA* or *phoA1* (**Fig. 1B**). Growth on PtdIn or GroPI was unaffected in the ΔM5 mutant and this strain also grew on the corresponding phosphomonoester substrate inositol 1-phosphate (IP1), but not the alternative product of GroPI hydrolysis, glycerol 3-phosphate (G3P) (**Fig. S2**). These data suggest *F. johnsoniae* possesses specialised pathways to utilise PtdIn and GroPI and that IP1 is produced as the monoester metabolite, likely as the result of PLC activity.

**Figure 1.**
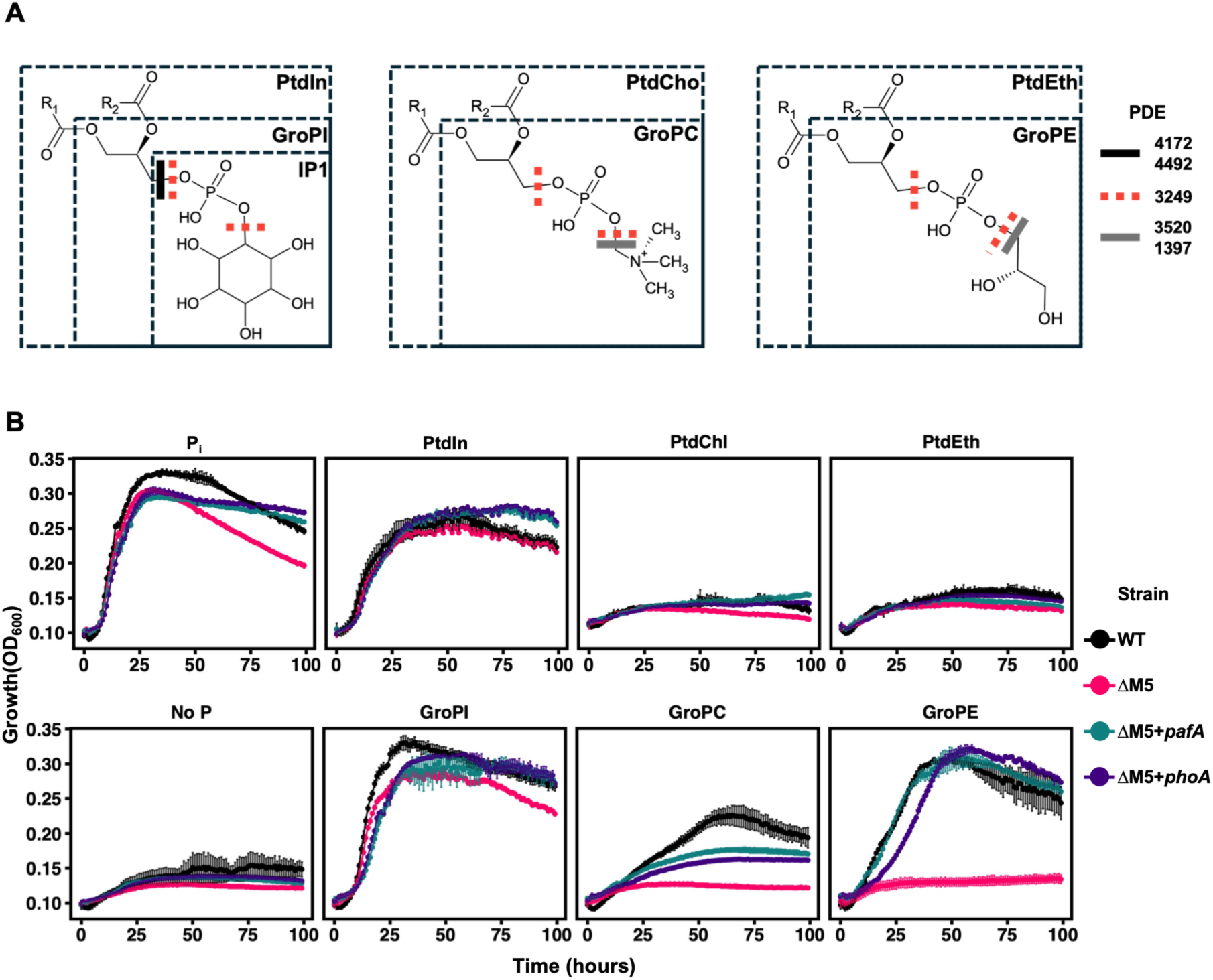
Growth of *Flavobacterium johnsoniae* on phospholipids and corresponding phosphodiester headgroups as a sole phosphorus (P) source. **(A)** Schematic of three common phospholipids tested in our study and the putative cleavage sites of the phosphodiesterases (PDEs) responsible for their breakdown in *F. johnsoniae*. Locus tags for putative PDEs are provided. **(B)** Wild-type (WT) *F. johnsoniae*, our previously characterised phosphomonoesterase ΔM5 mutant, and the ΔM5 mutant complemented with either *pafA* (*fjoh_0023*) or *phoA* (*fjoh_3249*) phosphomonoesterases grown on either PtdIn, PtdChl or PtdEth lipids or their corresponding soluble phosphodiester headgroups as a sole P source. Negative (no P) and positive (P_i_) controls are included. Results are the mean of triplicate cultures and error bars denote standard deviation. Data for all growth assays is representative of n=2 biological replicates.

### Pus1 and Pus2 mediate growth on PtdIn and GroPI

The genes encoding Pus1-4 are depicted in **Fig. 2A**. In addition to the PusCD transporter, *pus*1 encodes a putative PI-PLC (PI-PLC1, Fjoh_4172) and a putative calcineurin-like phosphatase (Fjoh_4173) that is structurally homologous to the recently characterised glycosylinositol phosphoceramide (GIPC)-PLC^39^, **Fig. S3**). Both proteins are predicted to possess Sec/SPI signal peptides (SignalP 6.0³⁹), indicating that they are co-translationally translocated into the periplasm. However, unlike previously characterised PI-PLCs, neither protein is predicted to be extracellular (DeepLocPro v1.0^40^) (**Fig. S4**). *pus2* encodes PhoA1, a multi-domain phosphatase (Fjoh_3249) containing a PhoA-like phosphomonoesterase domain (pf00249) and an uncharacterised glycerophosphoryl diester phosphodiesterase-like domain (pf13653, cd08577, PI-PLCc_GDPD_SF_unchar3). Both *pus* loci also harbour genes encoding extracytoplasmic function (ECF) sigma factor–anti-sigma factor pairs, which typically induce TBDT synthesis in response to substrate availability^41^. Pus3 and Pus4 possess distinct SLPs and contain numerous hypothetical lipoproteins making predictions about their function difficult.

**Figure 2.**
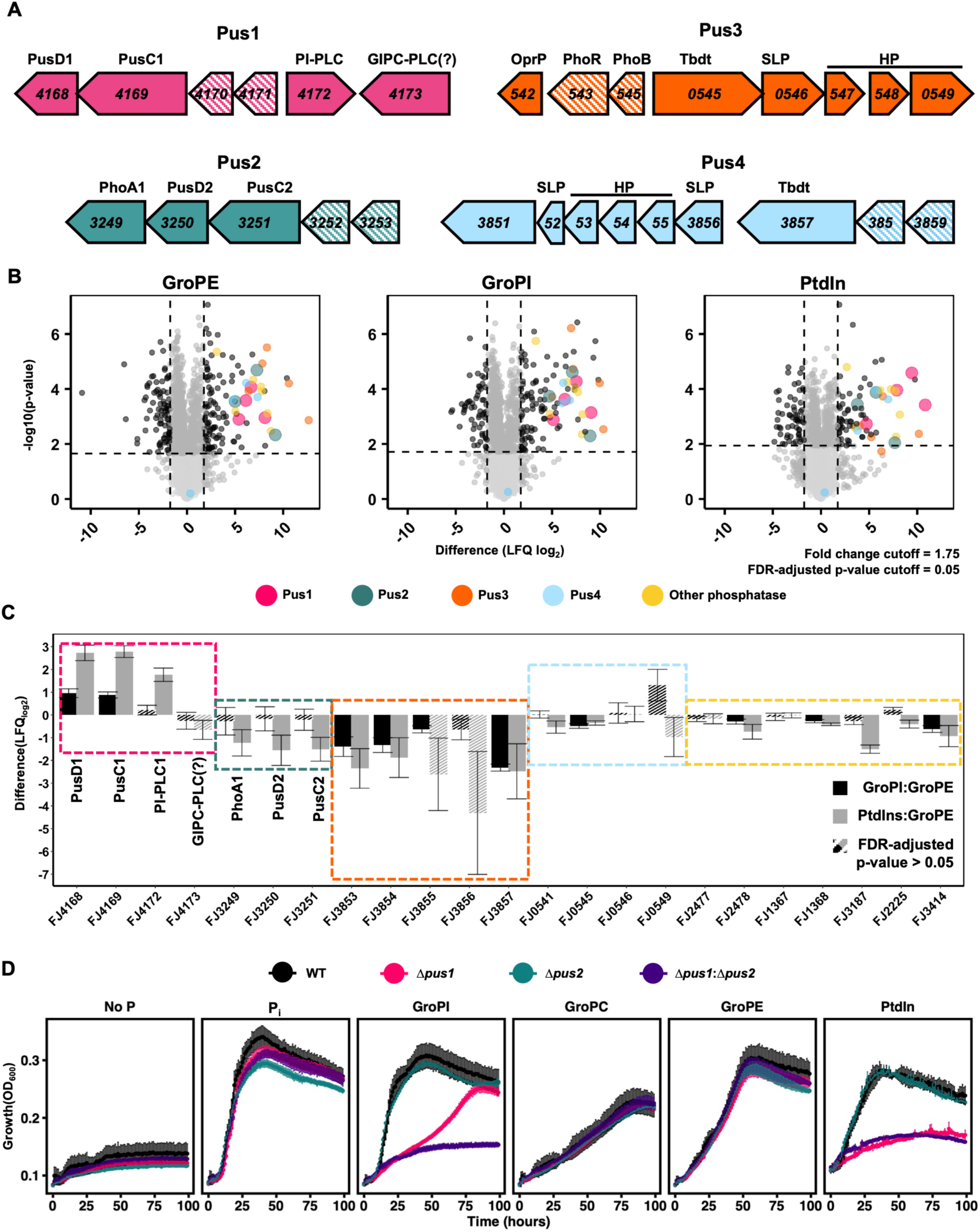
Contribution of Pus1 and Pus2 to growth on PtdIn and GroPI. **(A)** Organisation of the open reading frames (ORFs) in the Pus1–4 loci. Gene locus tags are shown; patterned ORFs denote regulatory elements. **(B)** Comparative proteomics of *F. johnsoniae* cultures (n = 3) grown on different P sources, showing differential protein levels relative to a phosphate-replete control. Proteins associated with Pus1-4 are highlighted in the indicated colours. Data show log_2_-transformed differences in label-free quantification (LFQ) values between treatment and control. **(C)** Relative abundance (log_2_ LFQ) of Pus-associated proteins and other phosphatases in cells grown on GroPI or PtdIn relative to GroPE. The mean of triplicate proteomes is presented with error bars depicting standard deviation. The Benjamini-Hochberg (BH) procedure was applied to control the False Discovery Rate (FDR) of P values across multiple comparison. **(D)** Growth of WT, Δ*pus1*, Δ*pus2* and Δ*pus1*:Δ*pus2* on various P sources. Lines show the mean of triplicate cultures and error bars represent standard deviation. Data is representative of n=4 biological replicates.

To test the hypothesis that Pus1 mediates growth on PtdIn, we performed comparative whole-cell proteomics of *F. johnsoniae* cells grown on PtdIn, GroPI or GroPE, alongside P_i_ -replete and P_i_ -limited controls. Relative to the P_i_ -replete control, growth on PtdIn, GroPI or GroPE was associated with elevated synthesis of proteins from Pus1, Pus2, Pus3 and Pus4, together with other proteins linked to organic P mobilisation (**Fig. 2B, Table S1**). Of these, however, only PusC1, PusD1 and PI-PLC1 reached their highest relative abundance during growth on PtdIn, indicating that Pus1 is the primary system mediating substrate capture and hydrolysis, and that it is subject to some degree of substrate-inducible synthesis, likely mediated by the ECF–sigma factor-anti-sigma factor regulator (**Fig. 2C**). The phylogenetically distinct PI-PLC2 (Fjoh_4492)^36^ encoded elsewhere in the genome was not detected under any condition.

We next deleted the Pus1 or Pus2 gene clusters individually (Δ*pus1* or Δ*pus2*) or in combination (Δ*pus1*:Δ*pus2*) and tested the ability of the resulting mutants to grow on different P sources (**Fig. 2D**). The single Δ*pus1* and Δ*pus2* strains and the double Δ*pus1*:Δ*pus2* mutant all grew like the wild type (WT) when P_i_, GroPC or GroPE was supplied as the P source. When GroPI or PtdIn was the sole P source, Δ*pus1* showed reduced growth relative to the WT, Δ*pus2* grew comparably to the WT, and Δ*pus1*:Δ*pus2* did not grow, indicating that Pus2 plays an auxiliary role in the absence of Pus1. Doubling the GroPI concentration from 50 µM to 100 µM reduced the growth defect of the Δ*pus1* mutant, suggesting that higher GroPI concentrations increase the efficiency with which Pus2 imports and hydrolyses the substrate and that GroPI is not the preferred substrate of PusCD2 (**Fig. S5**). Together, these data establish an essential function for the *pus1* and *pus2* in growth on GroPI and PtdIn, with *pus1* playing the major role.

### PusCD complexes are essential for growth on PtdIn and GroPI

To investigate the contribution of PusCD1 and PusCD2 to growth on GroPI and PtdIn, we generated in-frame deletions of the genes encoding each transporter. Only Δ*pusCD1* showed a reduced growth rate and final yield when grown on either GroPI or PtdIn, but grew like the WT on IP1 (**Fig. 3A**). Growth on GroPI and PtdIn was unaffected in Δ*pusCD2*; however, deleting *pusCD2* in the Δ*pusCD1* background to generate the double mutant Δ*pusCD1*:Δ*pusCD2* reduced growth relative to the Δ*pusCD1* mutant, confirming an auxiliary role for PusCD2. Complementation of Δ*pusCD1*:Δ*pusCD2* with *pusCD1* or *pusCD2* partially restored growth, confirming their function (**Fig. 3B**). The similarity between the full gene-cluster deletions (Δ*pus1* or Δ*pus1*:Δ*pus2*) and the transporter deletions (Δ*pusCD1* or Δ*pusCD1*:Δ*pusCD2*) supports the notion that, unlike previously characterised PI-PLCs that are secreted^42^, the phosphatases responsible for hydrolysing PtdIn and/or GroPI act in the periplasm. These data establish a key role for PusCD complexes in delivering organic P substrates across the OM, revealing a new function for TBDT–SLP complexes.

**Figure 3.**
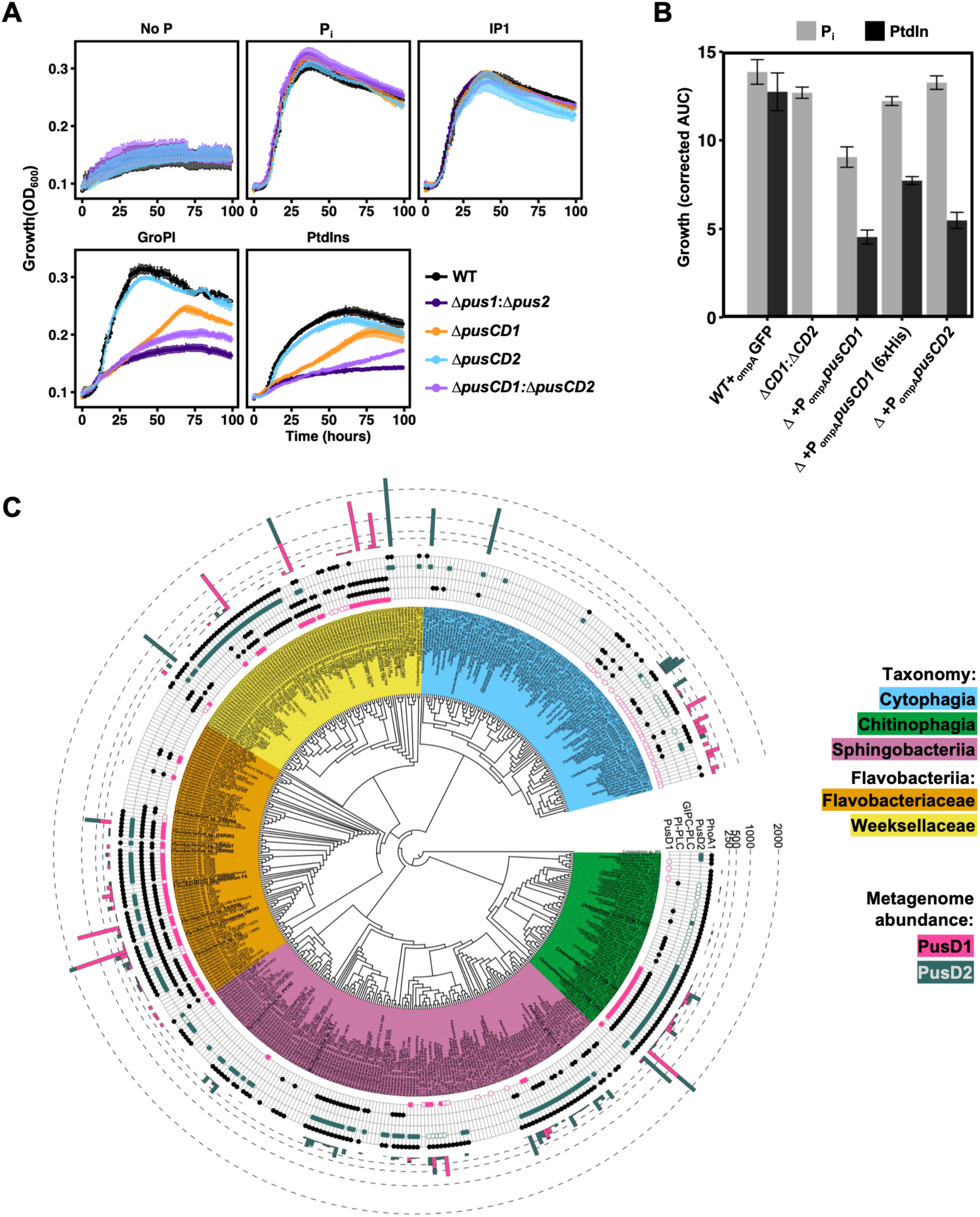
PusCD complexes mediate PtdIn and GroPI import. **(A)** Growth of the WT, Δ*pusCD1*, Δ*pusCD2* and Δ*pusCD1*:Δ*pusCD2* strains of *F. johnsoniae* on PtdIn or GroPI as the sole P source. **(B**) Growth of Δ*pusCD1*:Δ*pusCD2* complemented with either *pusCD1* or *pusCD2* under a constitutive (*ompA*) promoter. For each strain, the area under the curve (AUC) was corrected against residual growth in the no P control; the mean AUC of triplicate cultures is plotted, with error bars representing standard deviation. Data for all growth assays is representative of n=3 biological replicates. **(C)** Distribution of Pus components among terrestrial Bacteroidota genomes. Evolutionary relationships were inferred by maximum-likelihood multilocus sequence analysis (MLSA) based on three single-copy core gene nucleotide sequences (*guaA, gyrB, rpoB*). The presence or absence of Pus-like components is annotated in the inner rings as determined by BLASTP using the Pus1 and Pus2 proteins as queries. Hollow PusD squares represent PusD-like sequences which returned a BLASTP Expect-value of ≤1e^-40^ and >1e^-100^, with filled squares representing sequences with a value of ≤1e^-100^. The taxonomy of ORFs identified in plant-associated metagenomes, predominantly from rhizosphere soil, is shown in the outer ring; values represent the summed estimated gene abundance across all metagenomes, calculated in IMG/JGI from normalised scaffold coverage.

To assess the potential for Pus-mediated PtdIn utilisation in the rhizosphere, we screened terrestrial Bacteroidota genomes from the Integrated Microbial Genomes and Microbiomes (IMG/Joint Genome Institute) database for genes encoding Pus1 and Pus2 components (**Fig. 3C, Table S2**). Homologues were identified across the phylum, particularly among isolates from plant-associated environments. Using PusD1 and PusD2 as queries, we also screened plant-associated metagenomes^5^, confirming the presence of the transport machinery required to utilise phospho(inositol)lipids in Bacteroidota *in situ*. Plant-associated Bacteroidota isolates from our collection, including *Flavobacterium*, *Chitinophaga*, *Mucilaginibacter*, *Pedobacter*, and *Sphingobacterium* species, were screened for growth on PtdIn or GroPI as a sole P source. We found the presence of PusCD1 supported growth rates on these two organophosphate sources comparable to that on phosphate, whereas its absence resulted in slower growth despite the presence of other Pus1 and Pus2 components (**Fig. S6**).

### Cryo-EM structure of dimeric PusCD1 in complex with IP1

To gain insight into the substrate imported during growth on PtdIn, we purified PusCD1 from *F. johnsoniae* cells grown on a crude lipid extract (∼50% purity, Sigma-Merck) and determined the structure of the complex using cryo-EM. To aid purification, a Δ*pusCD1*:Δ*pusCD2* strain was transformed with a replicative plasmid encoding PusCD1 engineered to carry a C-terminal 6×His tag on PusC1. Correct synthesis, localisation and assembly of PusCD1 in the OM was confirmed by complementation of growth on PtdIn (Fig. 3B). As reported for glycan-binding SusCD-like complexes^18,43^, PusCD1 forms a dimer that interacts through the PusC components, with the ’pedal-bin’ PusD lids orientated 180° from one another (**Fig. 4A**). Global refinement of one distinct conformation (closed–closed, CC) resolved the structure of PusCD1 at 2.6 Å. The density map enabled rebuilding and refinement of a AlphaFold (V5)-predicted model (**Fig. 4B**). Each dimer of the CC complex contained extra density within an apparent ligand-binding cavity in PusD1 (**Fig. 4C**), consistent with the monoester product of PtdIn catabolism, IP1. Despite a large cavity formed by the PusC and PusD components (**Figs. S7** and **S8**) and our mutant growth data (**Fig. 3A**), no density corresponding to glycerol or the diacylglycerol lipid moieties of PtdIn was detected. In the CC state, each PusCD complex contained a single IP1 ligand within the solvent-excluded cavity, positioned 10.7 Å from plug domain loop (PL)1, but 3.8 Å and 3.1 Å from extended loop (EL)3 and EL10 of the barrel, respectively (**Fig. 4D**).

**Figure 4.**
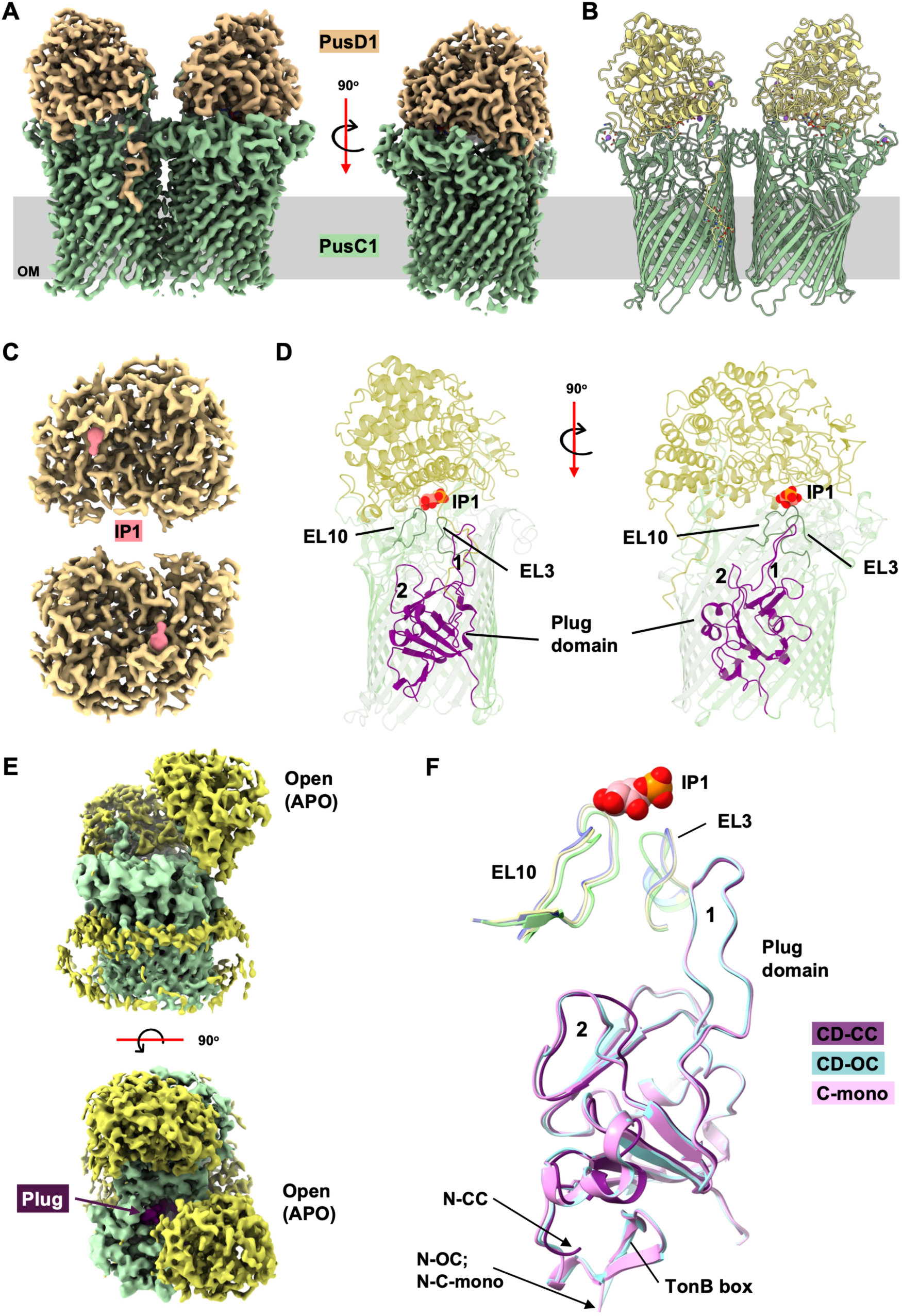
Structural insights into PusCD1-mediated substrate import during growth on PtdIn. **(A)** The cryo-EM density map of PusCD1 revealed the organisation of the dimeric structure with complexes in a closed-closed (CC) state and was used to generate the atomic model shown in **(B)**. **(C)** Each PusD1 contained unaccounted density consistent with the size and structure of IP1 (pink). **(D)** IP1 is observed bound in an apparent binding pocket of PusCD1 with no second ligand observed in the beta barrel. Extended loops (ELs) 3 and 10 of the PusC1 barrel along with loops 1 and 2 of the plug domain are highlighted. IP1 is shown as a space filled model. **(E)** The cryo-EM density map of the open-closed (OC) PusCD1 complex with one PusD1 lid open. From above, the plug domain (purple) inside the barrel is visible. **(F)** Comparison of the plug domain, EL10 and EL3 of PusC1 between the OC, CC and monomeric PusC1 (C-mono). EL10 and EL3 of PusC1 in the CC complex (green) have shifted relative to the OC (yellow) and C-mono (violet). The TonB box is absent in the CC state. Arrows indicating the N-terminus (N) of the polypeptide chain are highlighted.

### Ligand-bound PusD1 lid closure triggers periplasmic release of the TonB box

A subset of particles also yielded a ∼4 Å density map corresponding to an open–closed (OC) state, in which one PusD1 SLP was open with no detectable ligand bound (APO; **Figs. 4E** and **S9**). We independently purified PusC1 lacking PusD1 from *F. johnsoniae* grown on P_i_ as the sole P source and determined its structure to 2.3 Å (**Fig. S10**). No closed complexes without a bound ligand were detected in our dataset. Superimposition of the refined atomic models representing monomeric PusC1 and the OC and CC states of PusCD1 allowed us to determine the conformational changes in the barrel and plug domain of PusC1 induced by ligand binding and PusD1 ’lid’ closure (**Figs. 4F** and **5A**). Both the APO PusCD1 of the OC state and monomeric PusC1 contained density consistent with a TonB box (^203^EVLVVG^20^⁸) tucked inside the plug on the periplasmic side; this density was not detectable in PusC1 within the ligand-bound complex (**Figs. S9** and **S11**). Like the fructo-oligosaccharide (FOS) SusCD-like transport complex (SusCD^lev^)^43^, PusCD1 contains residues (Y209, Y594, F664) forming the ’aromatic lock’, which were shifted between the open and closed (ligand-bound) states^43,44^. However, unlike the FOS ligand in SusCD^lev^, upon lid closure the small IP1 ligand is unable to directly contact the plug domain while bound to PusD1 (**Figs. 4D** and **S12**). Consequently, PL1 does not differ between the OC and CC states, whereas ligand-induced shifts in density were detected for residues within EL3 (4 Å; S482), EL10 (1.5-3 Å) and PL2 (4-5 Å) (**Figs. 4F** and **S12**).

**Figure 5.**
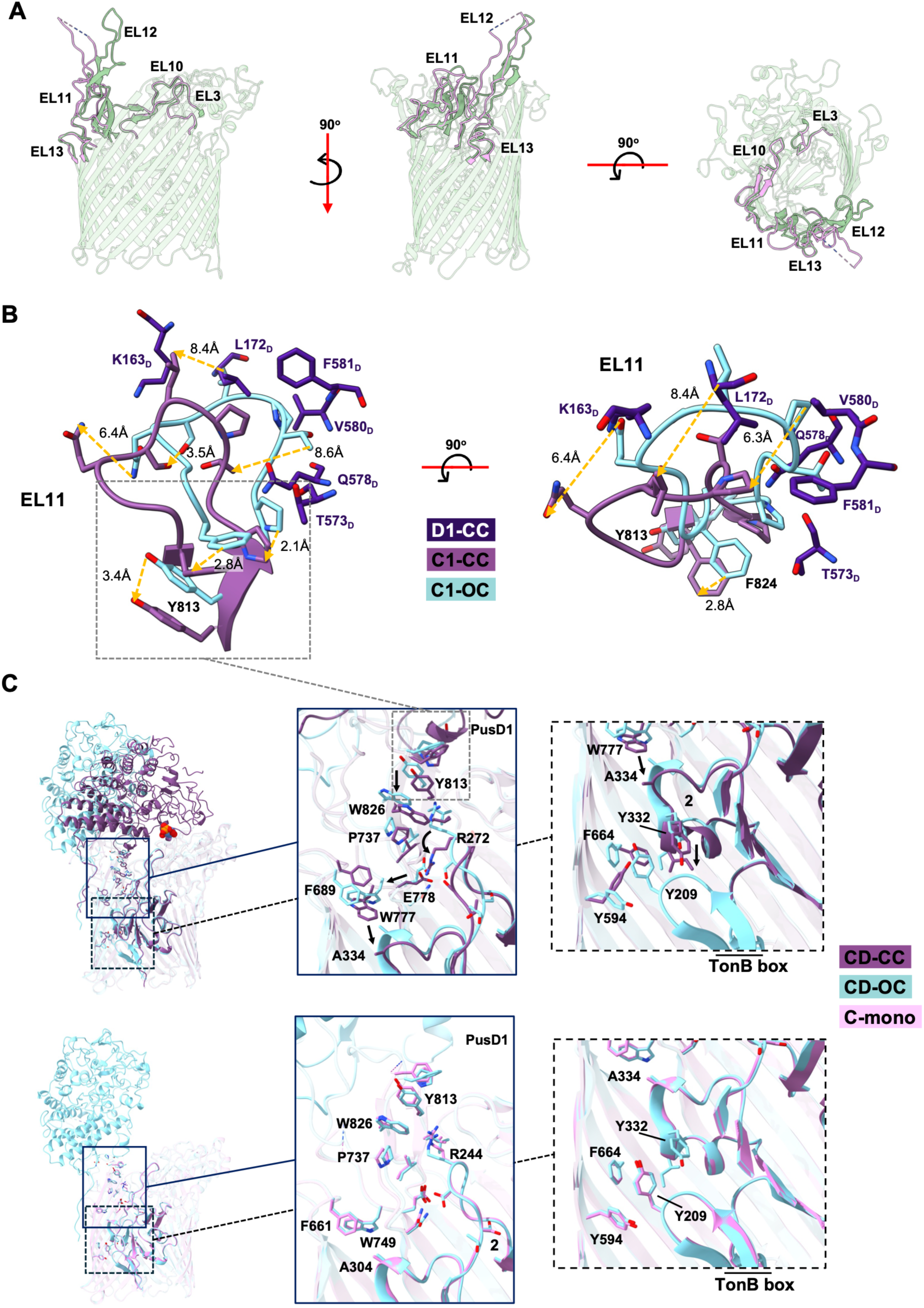
Conformational shifts in PusC1 extended loops in response to PusD1 lid closure. **(A)** PusC1 transmembrane domain visualised in the closed conformational position highlighting the extended loops (ELs) coordinating ligand sensing and signal transduction to the TonB box. **(B)** Superimposition of EL11 of PusC1 from the closed-closed (CC) and open-closed (OC) PusCD1 conformations demonstrating how the former is impacted by the proximity of PusD1 residues upon lid closure. The yellow dashed arrows show the shifted positions of key residues. IP1 is shown as a space-filling model. **(C)** Superimposition of the CC and OC PusCD1 complexes and monomeric PusC1 (C-mono) highlighting the similar aromatic lock regions of the PusC1 barrel and plug domains.

Conformational changes to EL11 and EL12 (major hinge loop) were also detected (**Fig. 5A**) We infer that PusD lid closure, involving the residues K163, L172_D_, T573_D_, Q578_D_, V580_D_, Q578_D_, F581_D_, initiates these changes by pushing on EL11 residues (P815, T817 P818, L819, Q821) causing several shifts between 6-9 Å, finishing with the downwards shift of Y785 (**Fig. 5B**). This shift transmits a signal down towards W777 (base of EL12) and F689 (EL13) via loop 2 of the plug (rotating R272), which pushes on A334 of the plug domain. This causes downward motion of Y332, breaking the triple aromatic lock (Y209, Y594, F664) and releasing the TonB box into the periplasm (**Fig. 5C**). 3D variability analysis (3DVA) of the OC state demonstrated mobility of the APO PusD1 lid, supporting a model where lid closure occurs spontaneously but is only stabilised by the presence of a bound ligand (**Movie S1**). This is supported by an apparent absence of closed PusCD1 complexes without a ligand. These data therefore emphasise the importance of both ligand-binding and the lid-closure mechanism in signalling periplasmic release of the TonB box, establishing a clear role for PusD1 contact in initiating a signal to the plug, particularly where a small ligand cannot directly interact with the plug domain upon binding.

### Ligand binding by PusCD1

The density maps readily resolved the residues in PusCD1 that bind IP1 (**Figs. 6A** and **6B**). In PusD1, R321, T323 and Y307 interact with the IP1 phosphate group, with R321 forming a salt bridge and T323 and Y307 making hydrogen bonds (**Fig. 6B**). R312 and R321 create a positive surface charge to attract the negatively charged phosphate group, a phenomenon not seen in the pocket of SusD^lev^ and BT2263, a SusD-like SLP isolated from *Bacteroides thetaiotaomicron* (*B. theta*) with a peptide ligand^18^ (**Figs. S7** and **S8**). The inositol moiety engages in π-stacking against the W85 side chain^45^ at the base of the shallow binding pocket, while D84 and D87 stabilise the hexose ring through interactions with the hydroxyl groups on carbons 2 and 5. In addition to these contacts with PusD1, hydroxyls on carbons 1, 4 and 6 of IP1 also form hydrogen bonds with PusC1 residues G884 and S886 on EL10, and with S482 on EL3; these ELs are shifted towards the substrate relative to the APO complex (**Fig. S12**). Using microscale thermophoresis (MST), we showed recombinant PusD1 binds GroPI with a relatively low affinity (*K*_d_ = 300-500 µM), somewhat lower than that (*K*_d_ ∼ 10^−4^ - 10^−5^ M) of the archetypal SusD for glycans (BT3701)^46^ (**Fig. 6C** and **S13)**. The π-stacking interaction provided by W85 explains why PusD1 has affinity for IP1, GroPI and the phosphorylated variants GroPI(4)P_1_, and GroPI(4,5)P_2_ but appears not to bind GroPE, and GroPC, which both lack the planar structure of an inositol ring to participate in this CH*-*π interaction (**Fig. 6C**). The product released by the putative GIPC-PLC, inositol phosphoglycan (IPG) also contains the IP1 moiety (**Fig. 6D**). Overlaying the IP1 moiety of IPG with IP1suggests potential room in the cavity between PusD1 and PusC1 to accommodate the additional sugars (**Fig. S14**).

**Figure 6.**
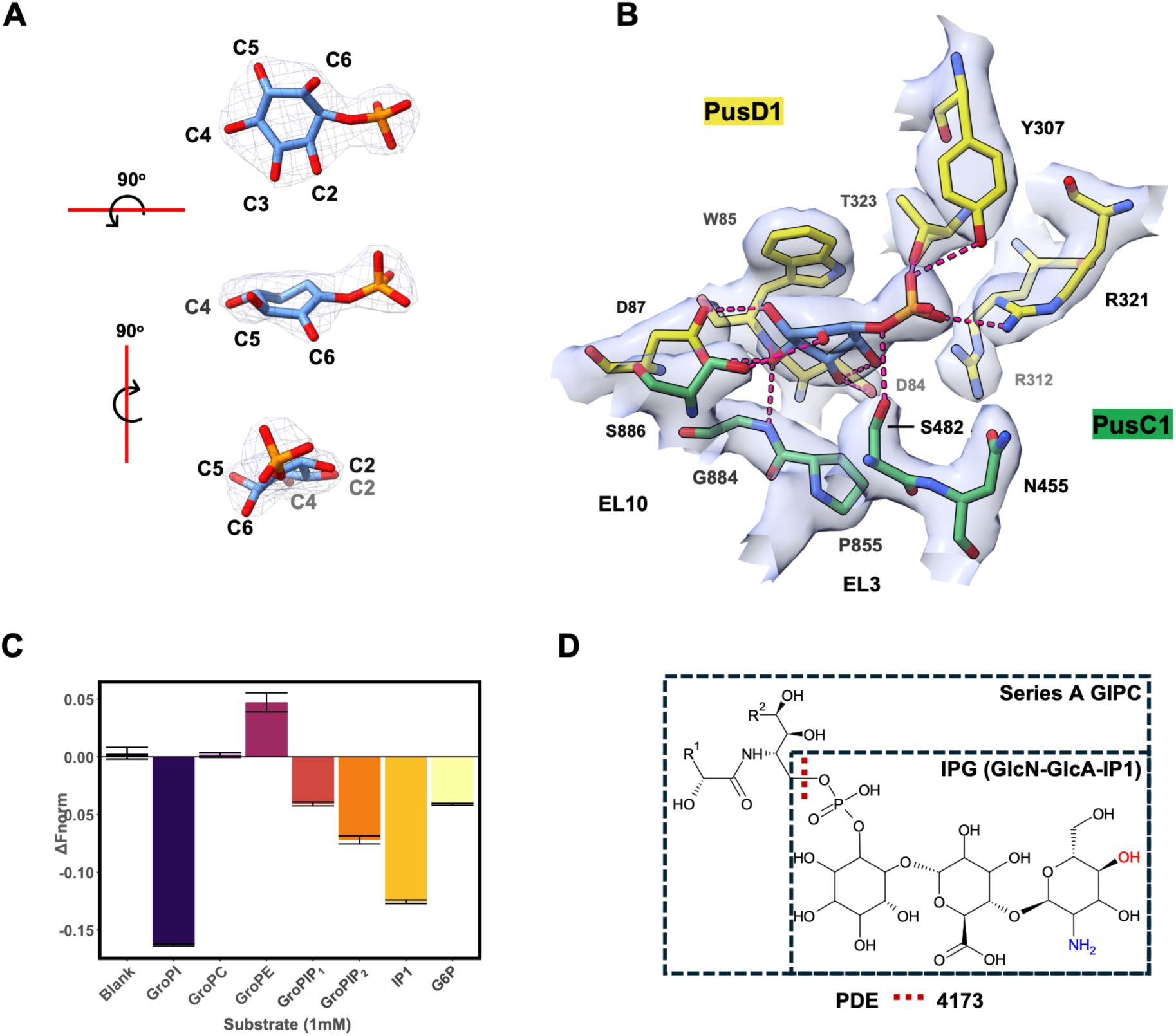
Structure-function relationships of ligand binding in PusD1. **(A)** Inositol phosphate (IP1) molecular structure depicting the numbered carbon on the inositol ring, with the corresponding density map overlaid (grey mesh). **(B)** IP1-PusD1 binding interactions overlaid with the density map (light blue). Predicted hydrogen bonds between the protein and IP1 are indicated by the red dashed lines. **(C)** Ligand-binding screen of recombinant PusD1 incubated against a range of phosphorylated substrates (1 mM) including GroPI and IP1 were performed by microscale thermophoresis (MST), using the magnitude of the shift as a proxy for the level of protein-ligand interaction (n=4). See **Fig. S13** for all substrates tested. **(D)** Example of a common Series A glycosylinositol phosphoceramide (GIPC) and the predicted site of hydrolysis by Fjoh_4173. GIPC glycosylation patterns vary by plant species and tissue type^47^.

### Convergent evolution of a common ligand-binding mechanism in PusD variants

Screening plant-associated Bacteroidota genomes (n=366, **Table S3**) showed that PusD1 and PusD2 are phylogenetically distinct, individually evolving from closely related SusD-like SLPs found in classical PUL associated with glycan utilisation, before radiating across diverse genera **(Fig. 7A)**. Across the Bacteroidota phylum, both *pusD1* and *pusD2* showed high genomic conservation, typically remaining associated with their corresponding *pus* loci. Unlike their SusD-like homologues, however, PusD ORFs were less frequently flanked (±6 kb) by CAZyme-encoding ORFs. PusD2 also has noticeable differences in its predicted size and fold relative to PusD1 (**Fig. 7B**), however PusD2 showed a similar affinity for GroPI (*K*_d_ = 200–400 µM) and a ligand-binding pattern comparable to that of PusD1 (**Fig. S13**), albeit with potential differences in relative affinity for IP1 and glucose-6-phosphate (G6P) (**Figs. 7C** and **S13**). In PusD1, the residues associated with binding IP1 and those adjacent in shallow pockets are conserved across PusD1 variants from diverse plant-associated Bacteroidota species, suggesting their importance in ligand binding (**Fig. 6D**). Using the same approach, we identified a similar region of conservation in the phylogenetically distinct PusD2, centred around a highly conserved Trp (W297) that may mediate π-stacking with the inositol ring of GroPI (**Fig. 6E**). Molecular docking of GroPI into both PusD1 and PusD2 (AutoDock Vina^48^) supported the hypothesis that these conserved regions in both proteins enable ligand binding via comparable mechanisms (**Figs. 6D** and **6E**). For PusD1 and PusD2, the GroPI poses shown had predicted affinity scores of -5.734 kcal/mol and -5.448 kcal/mol, respectively. Whilst both SLPs possess flanking Asp either side of the predicted Trp anchor and several Arg residues, PusD2 lacks the Tyr and Thr that are predicted to coordinate binding of the phosphate group within its pocket.

**Figure 7.**
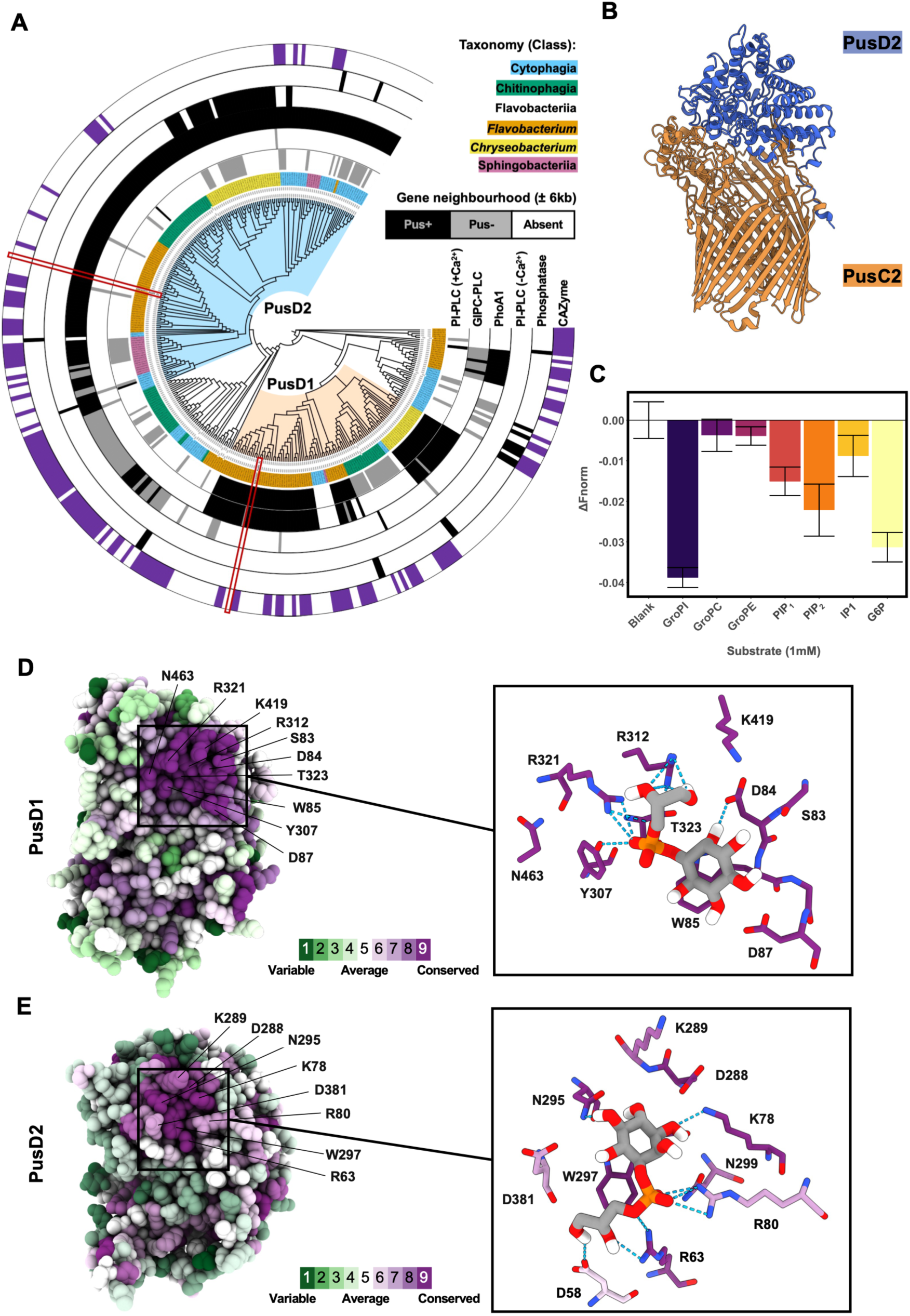
Convergent evolution of the binding mechanism in PusD homologs among plant Bacteroidota. **(A)** Phylogenetic reconstruction of PusD1 and PusD2 based on maximum likelihood in relation to their closest ‘SusD-like’ relatives, determined by BLASTP. Red boxes denote the *F. johnsoniae* PusD1 and PusD2 homologs. PusD1 (peach) and PusD2 (light blue) clades were determined by phylogenetic placement and genetic neighbourhood organisation. The outer ring shows the occurrence of CAZymes within 6 kb of ORFs encoding SusCD-like complexes. **(B)** AlphaFold 3 model of PusCD2 illustrating a similar predicted interaction between the PusD lid (blue) and the TBDT (PusC, orange) and the Tetratricopeptide Repeat (TPR) domains associated with a typical SusD-like SLP **(C)** Ligand-binding screen of recombinant PusD2 incubated against a range of phosphorylated substrates (1 mM) including GroPI and IP1 were performed using MST as per PusD1 in Fig. 6 (n=4). See **Fig. S13** for all substrates tested. (**D**) Residue conservation scores of AlphaFold 3 modelled PusD1 and PusD2 predicted by ConSurf^49^ focusing on the underside of the lid with predicted pocket resides highlighted. **(E)** Modelled interaction (Autodock Vina) between GroPI and either PusD1 or PusD2. Residue colouring is based on conservation as in panel D. For each ligand-protein interaction, the GroPI pose with the high-affinity score is shown. Hydrogen bonds (blue dashed lines) between ligand and residues are shown.

## Discussion

Plant-associated Bacteroidota are critical drivers of nutrient cycling^5,15,50^ and pathogen suppression^51–53^, making them prime targets for innovative microbiome engineering^9,20^. However, the molecular mechanisms governing their recruitment to the plant rhizosphere have remained largely elusive^9,20,36^. Building on our previous finding that Bacteroidota possess a specialised, energy-dependent transporter repertoire tailored for complex root exudates such as hemicelluloses^5,9^, we now report a distinct mechanism for host niche adaptation. Specifically, we show that *Flavobacterium* spp. utilise unique OM TBDT-SLP complexes to target plant PLs. This discovery redefines our understanding of these transport systems, which have traditionally been associated almost exclusively with complex carbohydrate import^4^. Our data also uncover a potential role for host PLs during root colonisation and plant-microbe interactions^27,39,54,55^, a hypothesis supported by comparative genomics showing that phospho(inositol)lipid utilization is a conserved, widespread trait across plant-associated Bacteroidota.

The root PM serves as the primary interface for host–microbe interactions^44^, with the sphingo(inositol)lipid GIPC comprising up to 40% of its outer leaflet^47,56^. At this boundary, glycero- and sphingo-(inositol)lipid turnover regulates both symbiotic signalling and plant–pathogen crosstalk^39,57,58^. The evolution of specialised gene clusters in Bacteroidota to exploit these abundant host lipids highlights a potential metabolic strategy enabling their rhizosphere and root endosphere colonisation across diverse plant species^9,20^. While our study confirms the utilisation of glycerophospho(inositol)lipids, the presence of a putative sphingolipid-acting phosphatase GIPC-PLC^39^ alongside PI-PLC in Pus1 suggests this cluster may also coordinate host GIPC hydrolysis.

Although PLs are likely abundant within the plant microbiome, how bacteria access them has remained poorly understood. The ecological rationale driving specialisation in phospho(inositol)lipid utilisation likely reflects a dual strategy for host immune evasion and P acquisition during rhizosphere and root endosphere colonisation^42^. Plant-associated Bacteroidota typically lack the high-affinity P_i_ and organophosphorus ABC transporters common to other dominant plant-associated bacterial taxa^7,19^. This genomic absence implies an evolutionary requirement for fundamentally distinct mechanisms to capture scarce forms of P within this host-associated niche, which we propose involves energy-dependent substrate import at the OM as opposed to the IM. Our data reveal Pus1 and Pus2 deliver a specialised capacity to utilise intact phospho(inositol)lipids, contrasting with an inability to utilise other common PLs such as PtdChl or PtdEth^56^. In addition to the presence of GIPC-PLC in the plant fungal pathogen *Botrytis cinerea*, the presence of PI-PLC homologs in diverse endophytic genera (including *Streptomyces*, *Pseudomonas*, and *Xanthomonas*)^35–37^ indicates a conserved role for phospho(inositol)lipid processing in successful root colonisation.

While PI-PLCs of other root endophytic strains are predicted to mimic the extracellular effector mechanisms seen in human pathogens^58^, the *Flavobacterium* PI-PLC1 and the putative GIPC-PLC are predicted to reside in the periplasm. Our genetic data support an essential role for the PusCD complex in mediating growth on PtdIn or GroPI, further substantiated by the capacity of the SLPs PusD1 and PusD2 to bind GroPI as well as IP1. In contrast, our cryo-EM data raise the possibility that IP1 may be released from PtdIn via extracellular PLC activity and subsequently imported by PusCD1. It is possible that extracellular release of PIPLC1 may have occurred from lysed cells in our overnight culture and supplementation with a second dose of PtdIn may have resulted in IP1 being released and captured by PusCD1. Higher protein synthesis of PI-PLC1 during growth on PtdIn does suggest this enzyme mediates lipid hydrolysis and further experimentation is required to determine the precise steps and localisation of utilisation in this bacterium.

PusCD1 and PusCD2 represent the first functionally characterised TBDT-SLP complexes dedicated to organophosphorus transport. Although both SLPs (PusD) are polyphyletic, independently evolving from closely related glycan-binding SLPs, structural modelling revealed a conserved mechanism for substrate binding; centring around a π-stacking interaction with a conserved tryptophan residue. The ability of PusD1 and PusD2 to bind IP1 and GroPI therefore likely represents an adaptation from glycans to phosphorylated sugars, which gives weight to the hypothesis that PusD1 may also bind IPGs released by the action of GIPC-PLC^39^. Unfortunately, experimental validation was not possible as GIPCs are not commercially available.

PusCD1 shares a common mechanism of signal transduction from surface-exposed ligand binding to TonB with that of recently characterised glycan-binding SusCD^lev42,43^. By utilising structural data on the monomeric PusC1, we show that PusD lid closure drives a trigger that cascades down the ELs of the barrel, to the aromatic lock. Unlike previous models derived from glycan-binding complexes, our data suggest that the signal for TonB box release does not require direct ligand-plug interaction, instead transmitting a signal cascade via: EL11 → plug loop 2 → EL12/3 → aromatic lock. Only when a ligand is bound does lid closure stabilise through interactions with EL3 and EL10 of the barrel. Here, EL11 acts as a sensor in addition to the hinge loop EL12. 3DVA supports this model whereby the flexible lid is only stabilised in the closed position long enough via the binding of the ligand to both subunits to emit a conformational change in the barrel and plug. Beyond *F. johnsoniae*, numerous phylogenetically distinct *pusCD*-like pairs are also present in the genomes of diverse plant-associated Bacteroidota. These loci frequently co-localise with alternative phosphatases, including phosphonate-hydrolysing genes, particularly those targeting aminophosphonate^19,59,60^. In addition, *pusCD*-like pairs are found adjacent to *glpQ*-like homologs in other plant- or gut-associated Bacteroidota. In these alternative functional contexts, a lack of a planar sugar ring comparable to inositol suggests the Trp-mediated stacking interaction is unlikely to occur, indicating an alternative mechanism of substrate interaction, which should be explored. Collectively, our study uncovers an overlooked family of organophosphorus transporters whose functional profiles likely diversify beyond a singular role in phospho(inositol)lipid import. Furthermore, the observed binding of recombinant PusD variants to GroPI(4)P_1_ and GroPI(4,5)P_2_ suggests a versatile capacity to interact with a broad range of phosphodiesters and perhaps even phosphotriesters. This molecular flexibility holds promising biotechnological implications for bioremediation applications targeting pesticides and sustainable P recovery from waste streams^61,62^.

## Materials and Methods

### Strains and growth conditions

*Flavobacterium johnsoniae* UW101 (DSM 2064), *Chitinophaga japonensis* DSM 13484*, C. pinensis* DSM 2588*, C. rupis* DSM 21039, and *C. terrae* DSM 23920 were purchased from the DSMZ collection. *Flavobacterium sp.* F52 was donated by the Cytryn Lab, Agricultural Research Organisation, Israel^63^. *Flavobacterium* sp. F6 and *Pedobacter* sp. A5 were donated by the Haubjerg Nicolaisen Lab, University of Copenhagen, Denmark ^64^. *Flavobacterium* spp. OSR003, OSR004, OSR005, OSR006, OSR007, OSR008, and *Sphingobacterium* sp. PV162 were previously isolated from the rhizosphere of *Brassica napus* L.^19^. *Mucilaginibacter* sp. IL1 was isolated from the roots (rhizoplane) of hydroponically grown lettuce. All Bacteroidota strains were maintained on casitone yeast extract (CY: 4g/L casitone, 1.25 g/L yeast extract, +/-2% (w/v) agar). For growth experiments investigating organophosphorus utilisation, Bacteroidota strains were grown in modified P-limited minimal A medium^65^, supplemented with 50 µM P source unless specified otherwise.

The organophosphorus substrates glucose 6-phosphate (G6P, CAS: 3671-99-6), glycerol 3-phosphate (G3P; CAS: 29849-82-9), glycerophosphorylcholine (GroPC, CAS: 28319-77-9), inositol 1,4,5-tris-phosphate (IP3; CAS: 141611-10-1), L-ɑ-phosphatidylinositol (PtdIn; from soybean; CAS: 97281-52-2) and phosphocholine (PC; CAS: 72556-74-2) were purchased from Sigma-Aldrich. Phosphatidylcholine (PtdChl; from soybean; CAS: 97281-47-5) and phosphatidylethanolamine (PtdEth; from soybean; CAS: 97281-51-1) were purchased from Larodan Research Grade Lipids. Glycerophosphoinositol (GroPI; CAS: 129830-95-1, glycerophosphoinositol 4-phosphate (GroPI(4)P_1_; CAS: 129830-96-2) and glycerophosphoinositol 4,5-bisphosphate (GroPI(4,5)P_2_; CAS: 111188-72-8) were purchased from Echelon Biosciences. Glycerophosphorylethanolamine (GroPE; CAS: 883288-78-6) was purchased from Cayman Chemical Company.

For growth assays, overnight CY-grown bacterial cultures were resuspended in minimal medium to serve as an inoculum (2% v/v). Assays were conducted at 28°C using 96-well microtiter plates and incubated in a Tecan Sunrise plate reader.

### Proteomic analysis of *F. johnsoniae*

*F. johnsoniae* cultures (n=3) were grown to mid-late growth phase in minimal medium supplemented with either 200 µM NaH_2_PO_4_ (‘low phosphate’), GroPI, GroPE or PtdIn, or with 1 mM NaH_2_PO_4,_ (‘high phosphate’). Cells were harvested by centrifugation at 4 °C at 8,000 x g for 10 mins and resuspended in 60 µL cold lysis buffer (10% w/v SDS, 100 mM TEAB in LC-MS grade water, pH 7.55) mL of cell culture followed by sonication and heating at 90 °C for 15 min. Cell debris was pelleted (16,000 x g at 4°C for 5 minutes) and the supernatant containing soluble protein was collected and quantified using the QuickStart^TM^ Bradford protein assay kit (BIO-RAD). Samples were prepared for mass spectrometry using an S-Trap Micro Kit (Protifi), combining tryptic digestion and on-column isolation of peptides. NanoLC-ESI-MS/MS was performed by the biOMICS Mass Spectrometry Facility, University of Sheffield, UK, on a ThermoScientific Orbitrap Exploris 480. The protein sequence database related to *F. johnsoniae* UW101 (UP000214645) was used for peptide identification using MaxQuant^66^ with quantification based on peak intensity of the precursor ion using LFQ with matched runs activated. Statistical analysis was performed using Perseus with significance values based on P values FDR-corrected using the Benjamini-Hochberg method^67^.

### Construction of *F. johnsoniae* mutants

The phosphomonoesterase mutant ΔM5 was created previously^38^. All mutants generated in this study followed the same allelic exchange protocol adapted from Zhu et al., 2017^68^. Regions flanking the gene of interest were cloned in into the suicide vector pYT313 carrying the *ermF* resistance gene and *sacB* counterselection marker and mobilised into the donor strain *E. coli* S17-1 *λ*pir. Allelic exchange was mediated by conjugation between donor and recipient. CY + 10% w/v sucrose was used as the counterselection medium to identify strains that had undergone a second recombination event. The generation of mutants was confirmed via PCR amplification using primers targeting 150 bp either side of the upstream and downstream homology arms and Sanger sequencing the amplicon (Azenta GENEWIZ). For complementation, *F. johnsoniae* genes were cloned downstream of the constitutive *ompA* gene promoter^5^ into the replicative plasmid pCP11. A full list of strains and primers used in this study can be found in **Table S4**. The pCP-derived, GFP-expressing plasmid, pAS43^69^, was provided by the Cytryn Lab, Agricultural Research Organisation, Israel.

### Comparative genomics

The Integrated Microbial Genomes and Microbiomes server (IMG/ Joint Genome Institute) was used to identify homologs of *F. johnsoniae* PusD1 (Fjoh_4168) and PusD2 (Fjoh_3250) in diverse plant-associated Bacteroidota genomes (**Tables S2 and S3**) using BLASTP (*E-*value cutoff: 1e^-40^). Sequences were aligned using MAFFT and clipped using ClipKIT ^70,71^. Phylogenetic analyses were performed with IQ-TREE2 using the parameters -m TEST -bb 1000 -alrt 1000^72^. Gene neighbourhoods of putative *pusCD1/pusCD2* orthologs were downloaded from IMG/JGI by exporting 6 kb regions upstream and downstream of PusCD ORFs. Gene neighbourhoods were annotated using Pyrodigal followed by HMMER and DBCan3 to identify phosphatases and CAzymes, respectively^73–76^. BLASTP against the IMG/JGI server was performed again to identify PUS1/PUS2-like phosphatases elsewhere in the Bacteroidota genomes, using the ORFs Fjoh_4172, Fjoh_4173, and Fjoh_3249 as query sequences. Annotated phylogenetic trees were created using Interactive Tree of Life (iTOL)^77^.

For constructing a phylogeny of Bacteroidota strains, BLASTN against the IMG/JGI server was performed to identify *guaA* and *rpoB* housekeeping gene sequences (*E-*value cutoff: 1e^-40^, 1 hit per query). Phylogenetic tree construction and annotation was then performed as above, with the additional step of concatenating the two sequences following alignment and clipping.

### Purification of Native PusCD1 complex

A 6-litre culture of *Flavobacterium johnsoniae* carrying the pCP11:p*ompA*:PusC1–C-terminal His₆-tag:PusD1 plasmid construct was grown overnight at 28 °C in minimal medium supplemented with 100 µM L-α-phosphatidylinositol (from soybean, SIGMA-MERCK). Cells were harvested at an OD_600_ of 0.25 by centrifugation at 5,000 × g for 20 min at 4°C. The pellet was resuspended in 30 mL of Buffer A (20 mM Tris pH 7.4, 300 mM NaCl), and a Complete EDTA-free protease inhibitor cocktail tablet was added to the resuspension before cells were lysed at a pressure of 10,000 psi using a pre-chilled French Pressure Cell Press. The lysate was centrifuged for 20 min at 20,000 x g at 4°C to remove cell debris, followed by ultracentrifugation at 204,252 x g for 1 hr at 4°C to isolate total cell membranes. The resulting pellet was resuspended and homogenised in 20 mL Buffer A containing 1.5% (v/v) N,N-Dimethyldodecylamine N-oxide (LDAO) using a glass Dounce homogenizer, then incubated for 1 hr at 4°C on a rotating platform. The sample was centrifuged again at 204,252 x g for 30 mins at 4°C and the supernatant containing solubilised membrane proteins was collected.

The protein complex was purified by Immobilised metal ion affinity chromatography (IMAC; PD-10, 2 mL Sepharose Fast Flow resin; Cytiva) where the sample was washed with 40 mL Binding Buffer 1 (25 mM HEPES, 1 M NaCl, 5 mM imidazole and 0.1% (w/v) Dodecyl-beta-D-maltoside DDM) followed by 25 mL Wash Buffer 1 (25 mM HEPES pH 7.4, 500 mM NaCl, 20 mM imidazole and 0.1% (w/v) DDM) and the protein complex was eluted using 12 mL Elution Buffer 1 (25 mM HEPES, 100 mM NaCl, 300 mM imidazole, and 0.05% (w/v) DDM). Elution fractions were pooled and concentrated to 5-8 mg/mL using a 100 kDa MWCO Amicon Ultra-4 centrifugal filter (Merck). The concentrated sample was further purified by size exclusion chromatography (SEC) on a Superdex 200 Increase 10/300 GL column (Cytiva) using a SEC Buffer 1 containing 10 mM HEPES, 100 mM NaCl and 0.05% (w/v) DDM. Fractions corresponding to the PusCD1 complex were determined via SDS-PAGE and collected, aliquoted, flash-frozen, and stored at -70 °C.

### Determination of the PusCD1 structure by cryo-EM

Purified PusCD1 at 8 mg mL⁻¹ was applied to glow-discharged Quantifoil R1.2/1.3 Cu 400 mesh grids. Grids were blotted and plunge-frozen in liquid ethane using a Leica vitrification device operated at 10 °C and 80% humidity. Cryo-EM data were collected on a Titan Krios transmission electron microscope operating at 300 kV and equipped with a SelectrisX energy filter (10 eV slit width) and a Falcon 4 direct electron detector. Images were recorded at a nominal magnification of 165,000×, corresponding to a calibrated pixel size of 0.74 Å (super resolution mode = 0.3645 Å). Movies were saved in TIFF format as 40-frame exposures with a total electron dose of 40 e⁻ Å⁻². Cryo-EM movies were analysed to generate 2D class averages and 3D reconstructions were performed using CryoSPARC^78^, with pipelines based on ^43,44,79^. Initial templates for picking were generated using the CRYOSPARC LIVE interface after ∼2000 images were initially processed and blob picking performed. After an initial 2D classification on the first 300 images, template picking using a box size 500 px was performed. In total ∼13,000 images were analysed, generating a total of 1.1M particles that were used for a 2-step 2D classification. Particle stack cleaning was performed using an *ab initio*-generated model, including several ‘junk’ classes and several rounds of heterogeneous refinement. A cleaned particle stack of ∼45k particles was subjected to initial non-uniform (NU) refinement to generate a refined volume for iterative 3D refinement. 3D classification (eight classes) was performed to identify heterogenous states. From this we captured two subsets of particles, reflecting a high-resolution CC state and the OC complex (two 3D classification rounds generated to OC-state stacks with 13.2k and 10.7k particles). All three particle stacks were subjected to particle re-extraction with an enlarged box size of 768 px down-sampled using Fourier cropping to 384 px. NU refinement of either state, followed by local refinement using a focused mask, was performed to increase the structural resolution of ligand binding and hinge domains. 3DVA on the OC state was performed to identify flexibility in the hinge mechanism responsible for mobility in the PusD1 lid domain. For detailed workflows, see **Figs. S15** and **S16** for the PusCD1 complex and PusC1 monomer, respectively.

### Production and purification of recombinant PusD homologs

Genes encoding PusD1 and PusD2 were PCR-amplified from *F. johnsoniae* gDNA, omitting their N-terminal signal peptide, and cloned into pET21a(+) in frame with the 6xHis tag. Expression plasmids were transformed into *E. coli* Rosetta(DE3)pLysS cells; Single colonies were grown overnight in LB with 100 µg/mL ampicillin and 34 µg/mL chloramphenicol and inoculated into 1 L of ZYM-5052 autoinduction medium to a final OD_600_ of 0.02^80^. Cultures were grown in the presence of antibiotic selection at 25 °C for 22 h with 150 rpm shaking. Cells were lysed via sonication, and pelleted at 20,000 g at 5 °C for 15 min. His-tagged PusD1 and PusD2 were purified using a three-stage procedure of IMAC, anion exchange, and SEC. IMAC was used for initial capture, using a 1 mL gravity column of Ni^2+^-chelated Sepharose Fast Flow resin (Cytiva). The sample was washed with 100 mL Binding Buffer 2 (25 mM HEPES pH 7.4, 1 M NaCl, 5 mM imidazole pH 8.0), followed by 100 mL Wash Buffer 2 (25 mM HEPES, 1 M NaCl, 50 mM imidazole), and step-eluted using Elution Buffer 2 (25 mM HEPES, 100 mM NaCl, 400 mM imidazole). An intermediate purification stage was required to further remove contaminating *E. coli* proteins; anion exchange chromatography was performed on a 1-mL Resource Q column (Cytiva) using a 50 mM Tris pH 7.4 buffer with a 40 mL gradient from 0 M to 500 mM NaCl. SDS-PAGE enabled determination of fractions with the lowest relative degree of contamination that were pooled and concentrated to 2 mL using a Vivaspin 30 kDa MWCO spin concentrator (Sartorius). SEC was performed with a Superdex 75 10/300 GL (Cytiva) using SEC buffer 2 (50 mM Tris pH 7.4, 50 mM NaCl, 10% glycerol). Purified aliquots were snap-frozen and stored at -70 °C prior to further use.

### Ligand binding assays using microscale thermophoresis (MST)

Recombinant PusD proteins were fluorescently labelled using a RED-tris-NTA 2^nd^ Generation His-tag labelling kit (Nanotemper). MST was performed using a Nanotemper Monolith NT.115 with excitation and MST power settings at 60% and 40%, respectively. Fluorescence shift was measured between 14-15 s post IR laser switched on. Initial binding checks were run using 1 mM substrate and 100 nM protein. For binding affinity estimates, 100 nM protein was titrated with decreasing GroPI concentration from 16 mM to 488 nM before MST. Dissociation constants were estimated using the *K*_d_ fit model within the Nanotemper MO.AffinityAnalysis software.

### Molecular docking of GroPI ligand into PusD SLPs

To predict the site and mechanism of GroPI-PusD interactions, the web-based platform of Autodock Vina (Webina), hosted by the Durrant lab was used. For PusD1, a real space model of the SLP was generated from our experimental data. For PusD2, the AlphaFold 3 (v6) model downloaded from the UniprotKB server was used (A5FEV7). 3D coordinates for GroPI were downloaded from PubChem. Autodock Vina assigns a random starting point for the ligand within each grid space provided. The following grid parameters were used: PusD1 - grid center: X -5 Y 16 Z 10, grid size : X 20 Y 12 Z 20, grid space : 0.375; PusD2 - grid center: X - 8 Y 16 Z 12, grid size : X 16 Y 14 Z 17.08, grid space : 0.375. To validate PusD1, a larger grid size (X 25 Y 18 Z 32) was used to cover a greater portion of the protein surface. Mode 1 showed the same poses and position of GroPI with comparable affinity (−5.759).

## Acknowledgements

This work was funded by grants from the Biotechnology and Biological Sciences Research Council (IL, BB/T009152/1) and the Royal Society (IL, URF\R1\221708: AH URF\R1\191548). The research of BvdB was supported by a Wellcome Trust Investigator award (214222/Z/18/Z). In addition, BvdB has received funding from the European Research Council (ERC) under the European Union’s Horizon Europe research and innovation programme (grant agreement No. 101201180). We thank Dr Indrajit Lahiri and Dr Svetomir Tzokov of the Sheffield Cryo-Electron Microscopy Facility (https://sheffield.ac.uk/cryo-em) for supporting grid preparation and screening. For high-resolution reconstruction of PusCD1 complex we acknowledge Diamond Light Source for access (session BI34172-30) and support (local technician, Éilís Bragginton) of the cryo-EM facilities at the UK national electron Bio-Imaging Centre (eBIC), proposal BI34172. The monomeric PusC1 structure was determined at The Astbury Centre for Structural Molecular Biology (University of Leeds) with support from Yehuda Halfon.

