## Supplementary Information for "Outer membrane ‘PusCD’ complexes coordinate phospho(inositol)lipid utilisation in plant Bacteroidota"

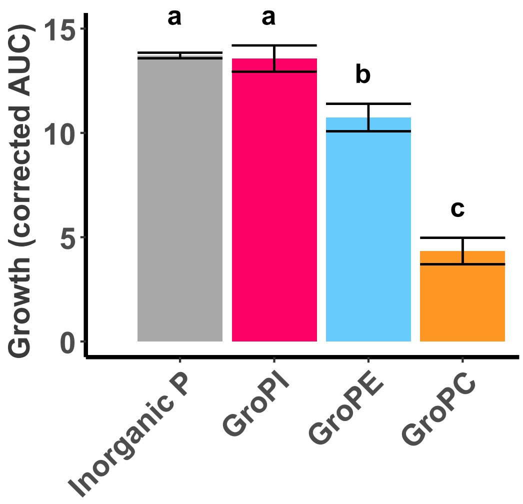


**Figure S1. Growth of wild-type *F. johnsoniae* on soluble headgroups of abundant glycerophospholipids.** Area under curve (AUC) values were calculated from growth curves shown in Fig. 1B. Analysis of variance was used to compare means between phosphorus treatments (ANOVA: F = 203.3, d.f. = 4, 10). Means that do not share a letter are significantly different by Tukey HSD test at p<0.001. Error bars represent standard deviation of triplicate cultures. Data representative of n=2 biological replicates.

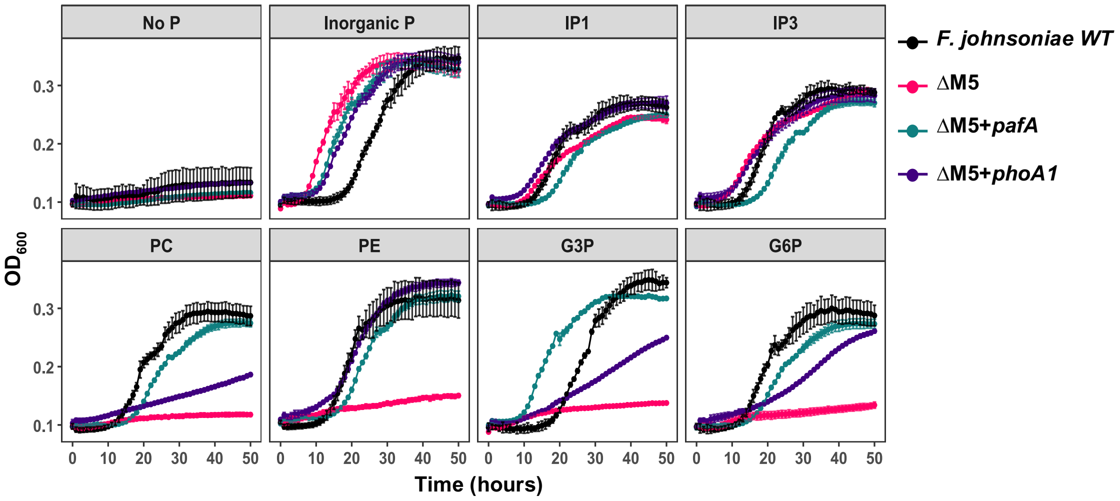


**Figure S2. Growth of *F. johnsoniae* on various organophosphorus substrates as a sole P source.** Growth of *F. johnsoniae* wild-type (WT), our previously characterised phosphomonoesterase-null mutant (ΔM5)[1], and the ΔM5 mutant complemented with either *pafA* (*fjoh_0023*) or *phoA1* (*fjoh_3249*) alkaline phosphomonoesterases on phospholipid monoester headgroups. Abbreviations: IP1, inositol 1-phosphate; IP3, inositol triphosphate PC, phosphocholine; PE, phosphoethanolamine; G3P, glycerol 3-phosphate; G6P, glucose 6-phosphate. Data representative of n=2 biological replicates.

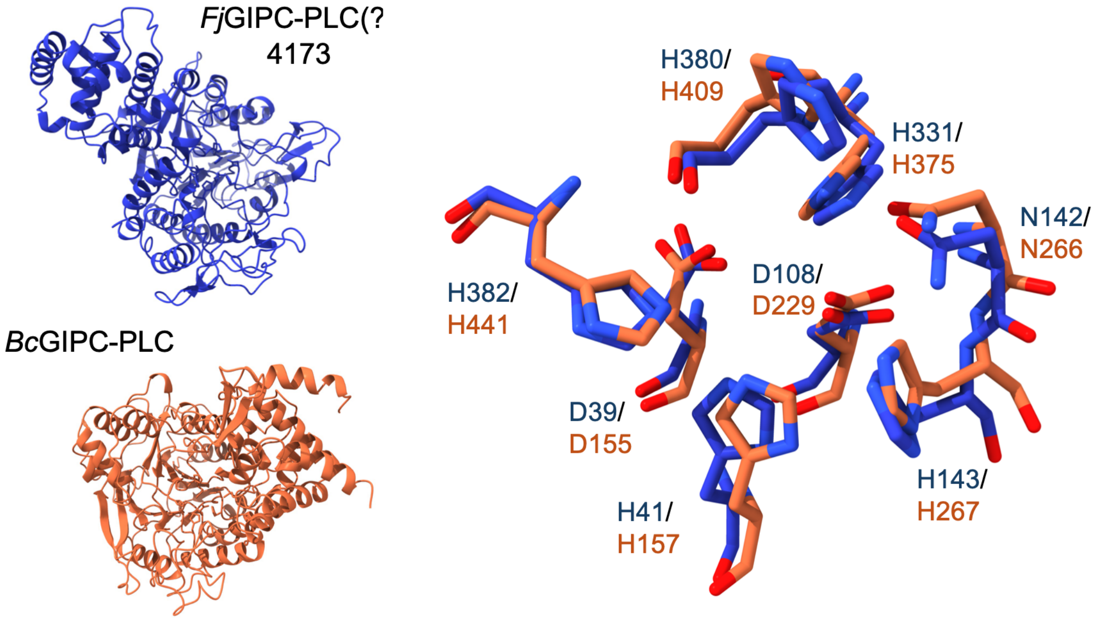


**Figure S3. Structural comparison of GIPC-PLCs.** AlphaFold 3 (v6) models of *Fj*GIPC-PLC (Fjoh_4173, blue) and the recently characterised *Bc*GIPC-PLC1 (BCIN07g04350, orange)[2], demonstrating conserved amino acid residues in the catalytic core where hydrolysis of GIPC into ceramide and inositol phosphoglycan is predicted to occur.

**SignalP 6.0**

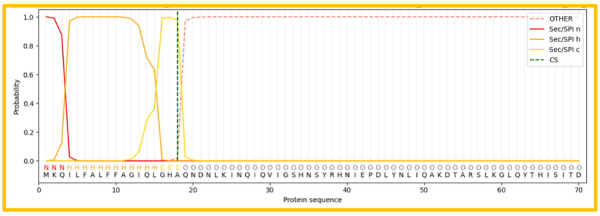


| Protein type | Other | SP(Sec/SPI) | LIPO(Sec/SPII) | TAT(Tat/SPI) | TATLIPO(Tat/SPII) | PILIN(Sec/SPIII) | Cleavage site |
|---|---|---|---|---|---|---|---|
| **Probability** | 0.000228 | 0.99909 | 0.000194 | 0.000187 | 0.000151 | 0.000147 | pos:18-19 0.9776 |

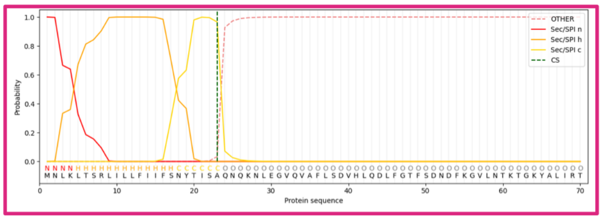


| Protein type | Other | SP(Sec/SPI) | LIPO(Sec/SPII) | TAT(Tat/SPI) | TATLIPO (Tat/SPII) | PILIN(Sec/SPIII) | Cleavage site |
|---|---|---|---|---|---|---|---|
| **Probability** | 0.000279 | 0.998988 | 0.000235 | 0.000209 | 0.00017 | 0.000165 | pos:23-24 0.9657 |

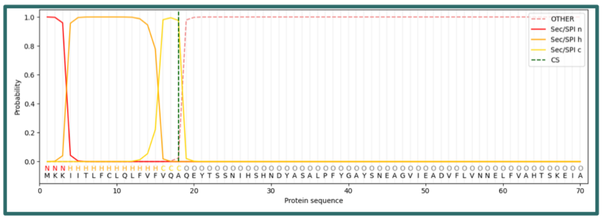


| Protein type | Other | SP(Sec/SPI) | LIPO(Sec/SPII) | TAT(Tat/SPI) | TATLIPO (Tat/SPII) | PILIN(Sec/SPIII) | Cleavage site |
|---|---|---|---|---|---|---|---|
| **Probability** | 0.000186 | 0.999201 | 0.000181 | 0.000167 | 0.000135 | 0.000139 | pos:18-19 0.9760 |

**DeepLocPro 1.0**

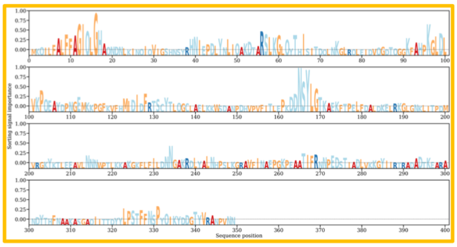


| Localisation | Cell wall & surface | Extracellular | Cytoplasmic | Cytoplasmic membrane | Outer membrane | Periplasmic |
|---|---|---|---|---|---|---|
| **Probability** | 0.0021 | 0.2525 | 0.0812 | 0.003 | 0.0837 | 0.5775 |

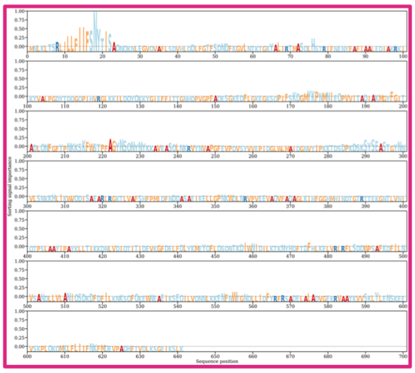


| Localisation | Cell wall & surface | Extracellular | Cytoplasmic | Cytoplasmic membrane | Outer membrane | Periplasmic |
|---|---|---|---|---|---|---|
| **Probability** | 0.0013 | 0.1465 | 0.008 | 0.084 | 0.2425 | 0.5176 |

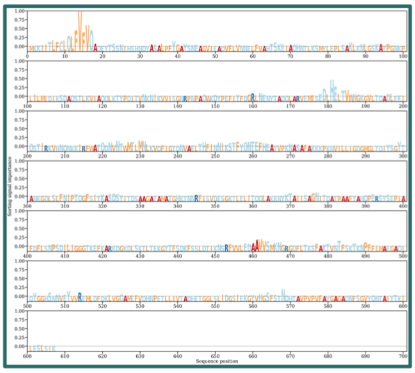


| Localisation | Cell wall & surface | Extracellular | Cytoplasmic | Cytoplasmic membrane | Outer membrane | Periplasmic |
|---|---|---|---|---|---|---|
| **Probability** | 0.0023 | 0.2277 | 0.0057 | 0.0355 | 0.1291 | 0.5998 |

**Pus-associated phosphatase**

***Fj*PI-PLC**

***Fj*GIPC-PLC**

***Fj*PhoA1**

**Figure S4. Predicted cellular localisation of Pus-associated phosphodiesterases (phosphatases).** Amino acid sequences of *Fj*PI-PLC (Fjoh_4172, orange), *Fj*GIPC-PLC (Fjoh_4173, pink) and *Fj*PhoA1 (Fjoh_3229, green) were used as queries into the bioinformatics programs DeepLocPro 1.0 and SignalP 6.0 to determine their most likely site of localisation and the mechanism for secretion[3,4]. According to SignalP, all three possess SPI cleavage sites which would be consistent with secretion to the periplasm— the predicted localisation by DeepLocPro.

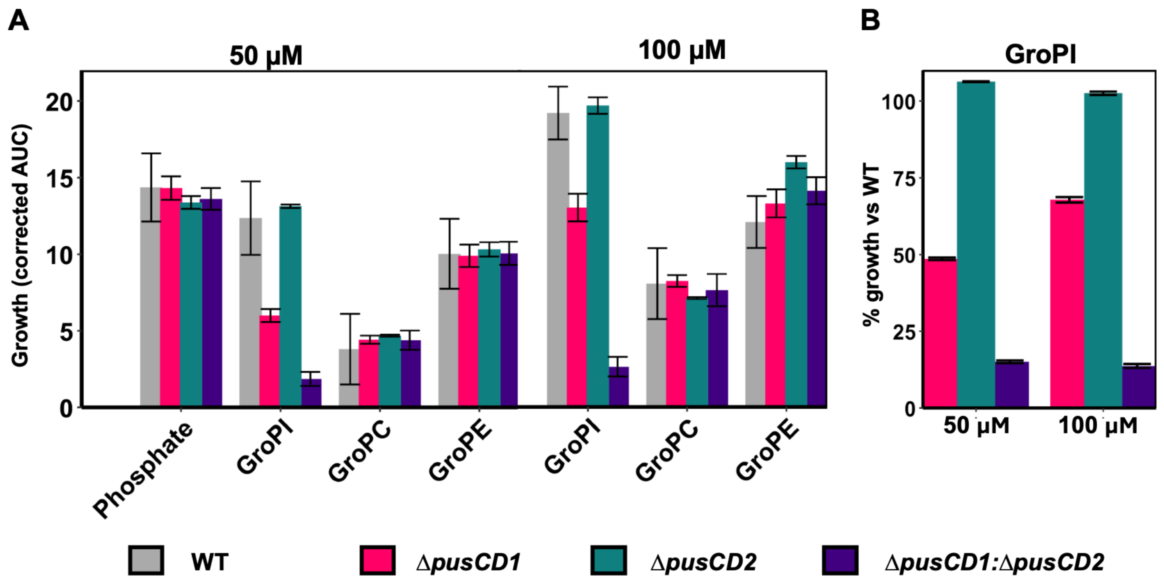


**Figure S5. Impact of substrate concentration on the efficiency of *F. johnsoniae* strains to grow on phosphodiester headgroups as a sole phosphorus source. (A)** Growth was quantified using the sum "Area under curve" (AUC) for Pus cluster mutants and the parental wild-type grown with either 50 µM or 100 µM glycerophosphoinositol (GroPI), glycerophosphoethanolamine (GroPE) or glycerophosphocholine (GroPC). AUC values have been corrected by deducting the AUC of a no phosphate negative control from each condition. Upon doubling the concentration of GroPI, the relative impact of deleting *pus1* decreased. **(B)** To highlight the decreased impact of mutation at the higher substrate concentration, the proportional difference in AUC was plotted for GroPI-grown cultures. Values represent the mean of triplicate cultures. Error bars denote standard deviation. Data presented is representative of n = 3 biological replicates.

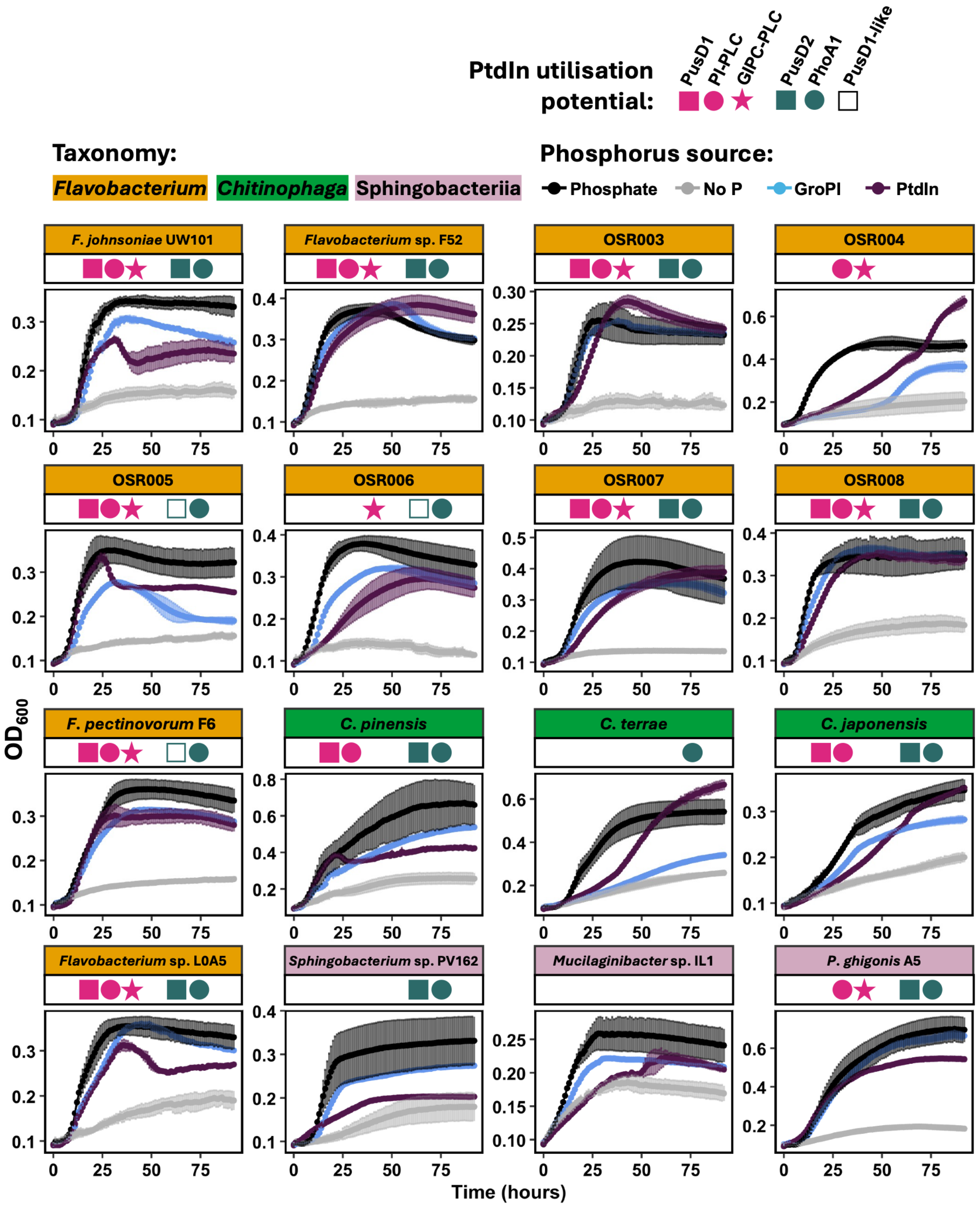


**Figure S6. Phospho(inositol)lipid utilisation in plant-associated Bacteroidota.** Growth screen of plant-associated Bacteroidota isolates on 50 µM GroPI and PtdIn. Symbols above each growth curve represent the respective strain's homology to the Pus components in *F. johnsoniae* DSM2064 as determined by BLASTP. Sequences with hits against PusD1 with 'e-values of >1e-100 and <1e-140 were designated 'PusD1-like', due to falling outside of the PusD1 clade identified in Fig. 6a of the main text, as well as being found within distinct gene neighbourhoods to Pus1.

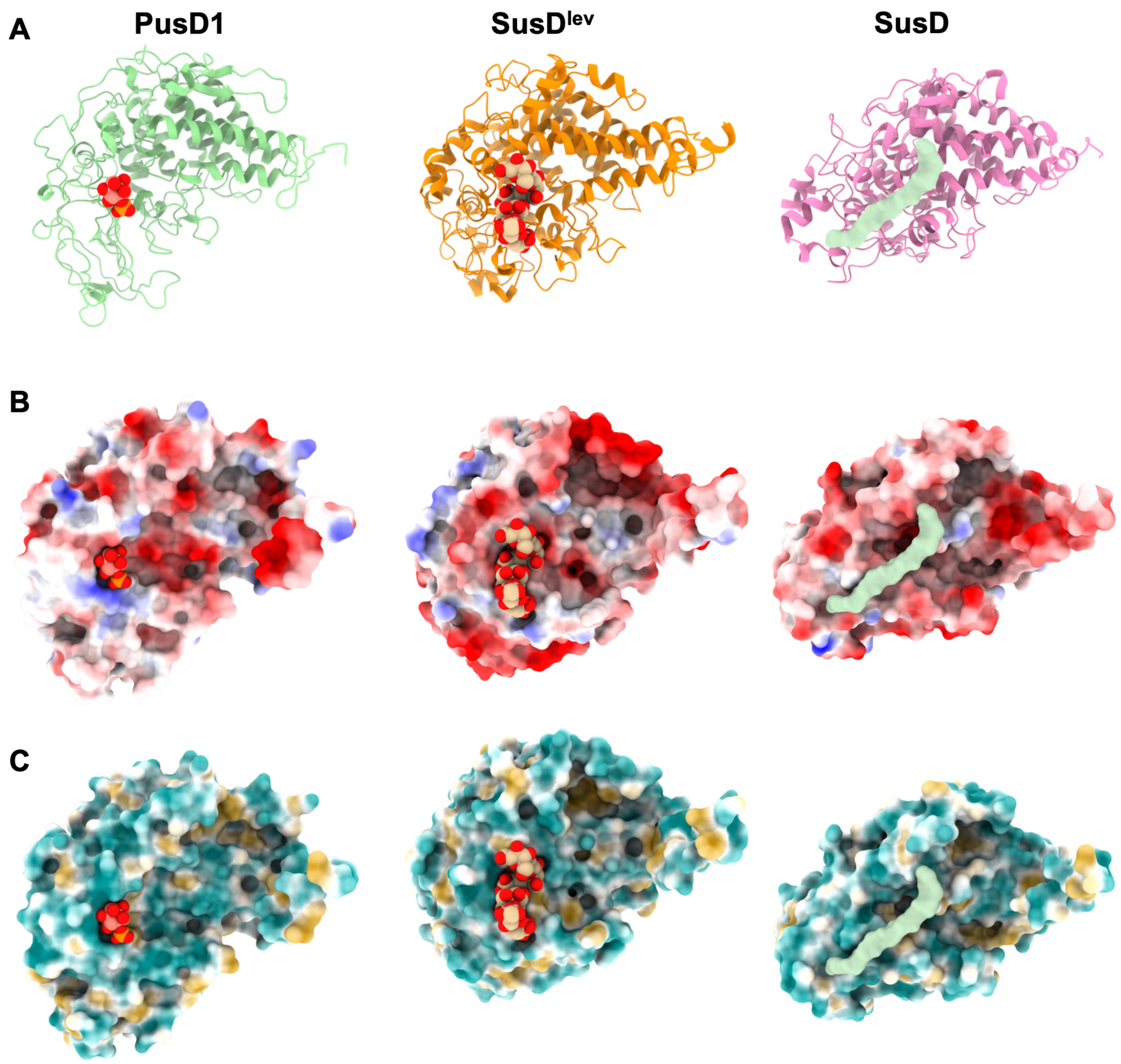


**Figure S7. Comparison of PusD1 with previously characterised SusD surface-exposed lipoproteins (SLPs).** PusD1 (Fjoh_4168) is bound to the inositol 1-phosphate ligand (this study), SusD$^{lev}$ (BT1763) is bound to fructo-oligosaccharide and SusD (BT2263) is bound to an unidentified linear peptide, modeled as deca-glycine[5,6]. All ligands are presented as space-filling models. **(A)** Cartoon model depicting the backbone features showing the Tetratricopeptide Repeat (TPR) domains common to all SusD-like lids. **(B)** Modelled surface electrostatic potential revealing a region of positive charge (blue) in PusD1 that attracts the negatively charged phosphate group and the smaller binding cavity compared to the other SusD proteins. **(C)** Modelled surface hydrophobicity of all three SLPs. Hydrophilic regions are blue and hydrophobic regions are gold.

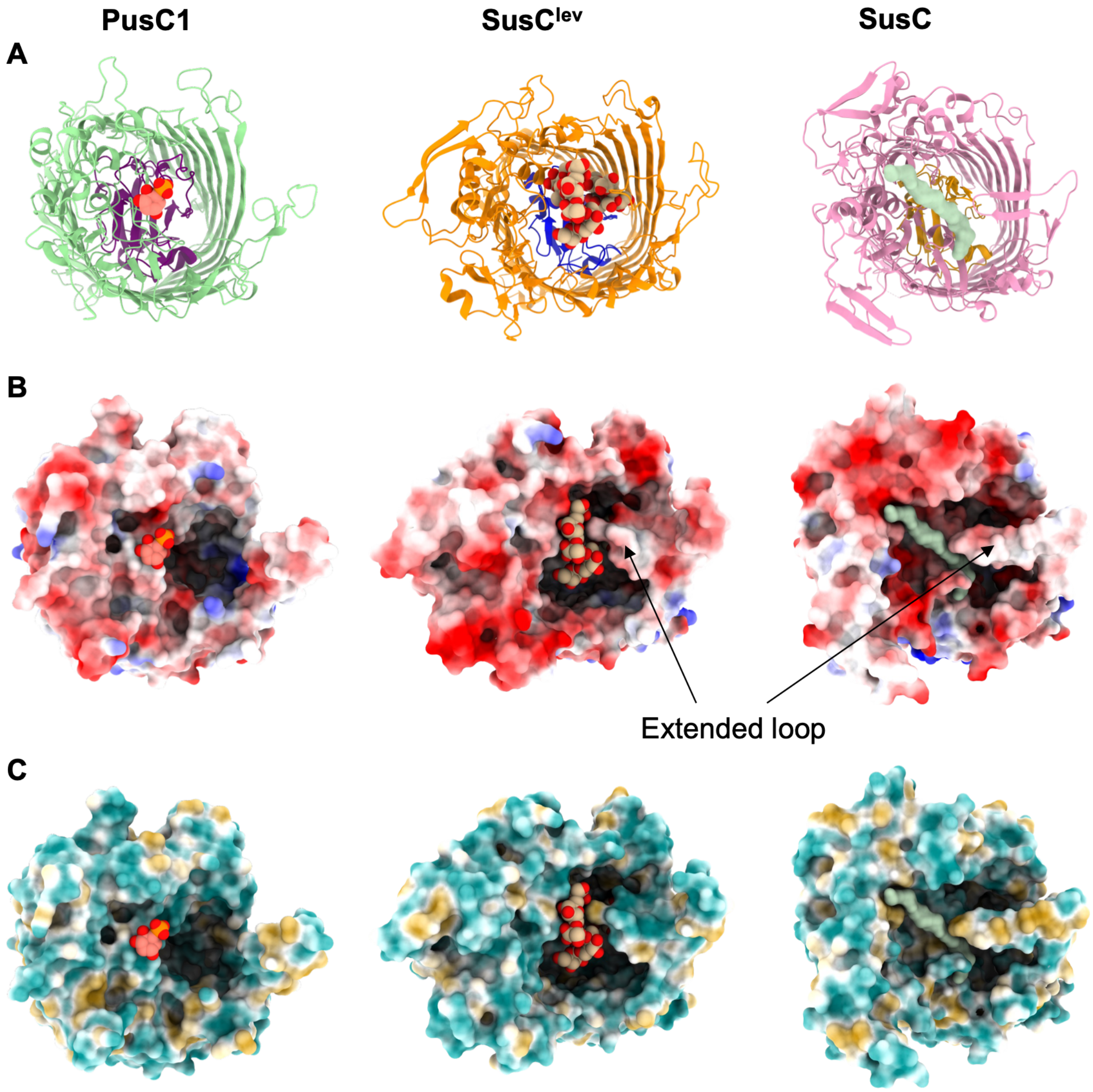


**Figure S8. Comparison of PusC1 with previously characterised SusC proteins.** PusC1 (Fjoh_4169) is bound to the inositol 1-phosphate (IP1) ligand, $SusC^{lev}$ (BT1762) is bound to fructo-oligosaccharide (FOS) and SusC (BT2264) is bound to an unidentified peptide, modelled as deca-glycine[5,6]. All ligands are presented as space-filling models. **(A)** Cartoon model depicting the backbone features, distinguishing the transmembrane beta-barrel and plug domains of the transmembrane domains. **(B)** Modelled surface electrostatic potential showing the presence of a horizontal extended loop on the barrel of each glycan-importing SusC that is absent from PusC1. Positively charged regions are blue and negatively charged regions are red. **(C)** Modelled surface hydrophobicity of the three SLPs. Hydrophilic regions are blue and hydrophobic regions are gold.

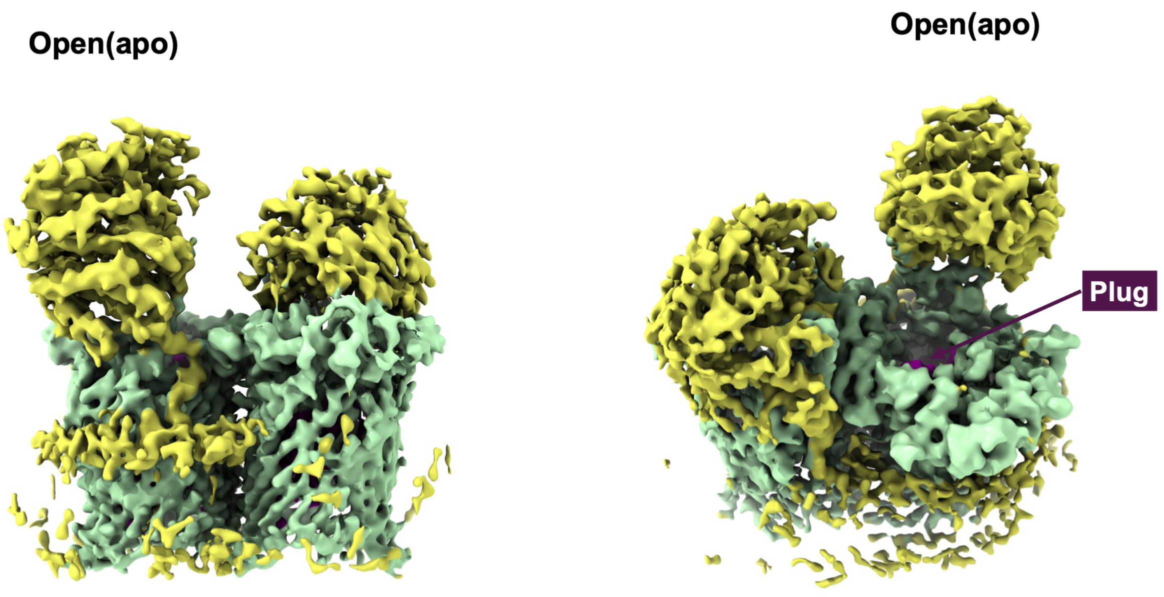


**Figure S9. 3D reconstruction of PusCD1 in the open-closed (OC) state.** Two alternative angles of the complex are shown in addition to Fig. 4E. The plug domain (purple) inside the barrel of the open (APO) PusCD1 complex is visible.

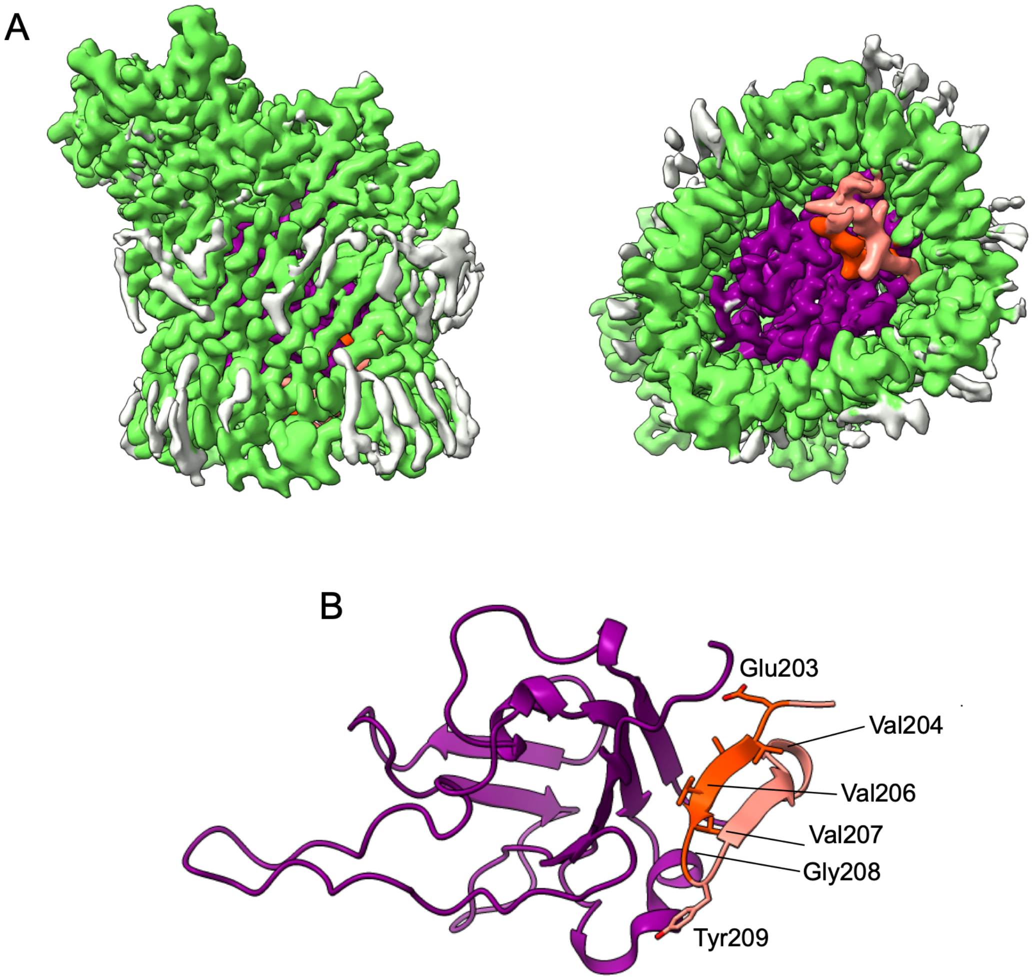


**Figure S10. 3D reconstruction of monomeric PusC1 lacking the PusD1 lid. (A)** Density map of PusC1 resolved to 2.3 Å. **(B)** Cartoon model of the plug domain of the monomeric PusC1 with the TonB box (coloured salmon) inaccessible to periplasmic TonB. TonB box residues are labelled. The conformation of the plug domain of monomeric PusC1 was identical to that of the open-state PusCD1, indicating PusD1 closure is an important trigger for TonB box release.

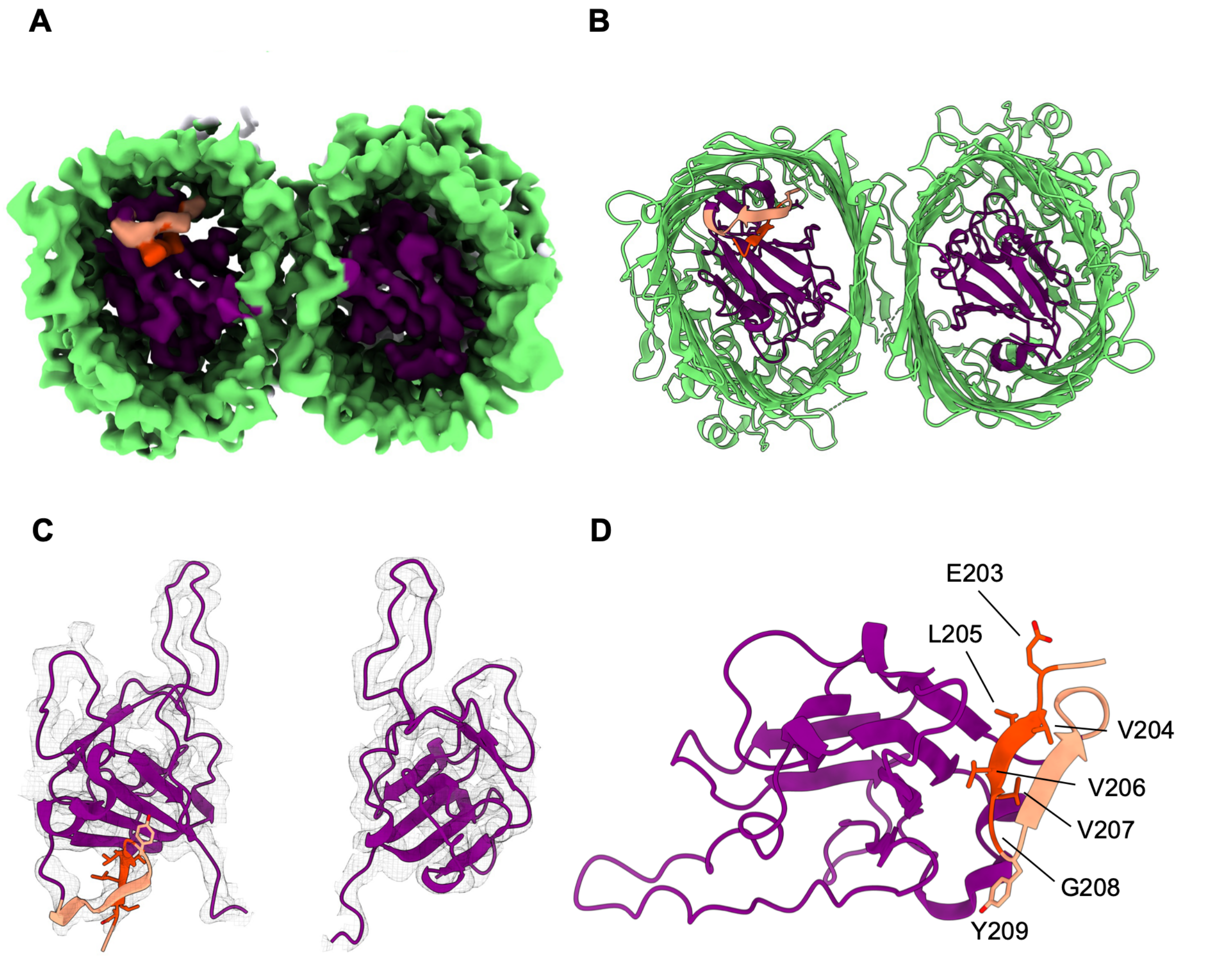


**Figure S11. Positioning of the TonB box in the open-closed (O-C) state of the PusCD1 complex. (A)** Bottom-view of density maps representing both PusC1 barrels (green) occupied with their corresponding plug domains (purple). The left barrel, corresponding to the open state, contains extra density (light salmon) including the TonB box (orange). **(B)** Cartoon model of both PusC1 barrels based on the observed density map **(C)** The N-terminal region of the PusC1 plug domain, containing the predicted TonB box (black) with side chains represented ($^{203}$EVLVVG$^{208}$) for the TonB box) and density overlaid (black mesh). **(D)** Superimposition of the plug domains of PusC1 in the closed- (light green) and the open-state (purple) maps. The extra density corresponding to the region containing the TonB box (orange) is coloured peach. Side chains of the TonB box are shown.

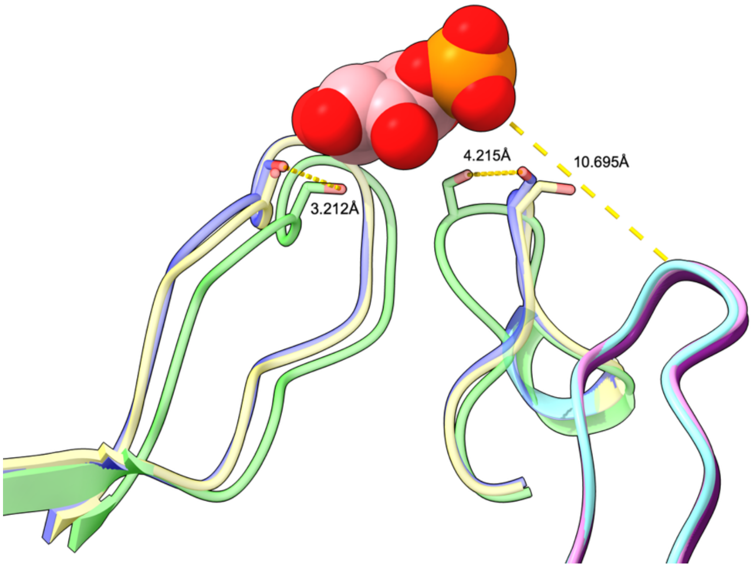


**Figure S12. Unambiguous shifts of extended loop (EL)3 and EL10 of the PusC1 TBDT.** The distance (angstroms, Å) of the conformational shifts between side chains groups of S886 (EL10), S482 (EL3) in the open complex (pale yellow) or monomeric PusC1 (pale blue) versus the closed complex (light green) are provided. The distance (Å) between the oxygen on the phosphate groups of inositol 1-phosphate (IP1) and G299 of the plug loop 1 is also provided. IP1 is shown as a space-filled model.

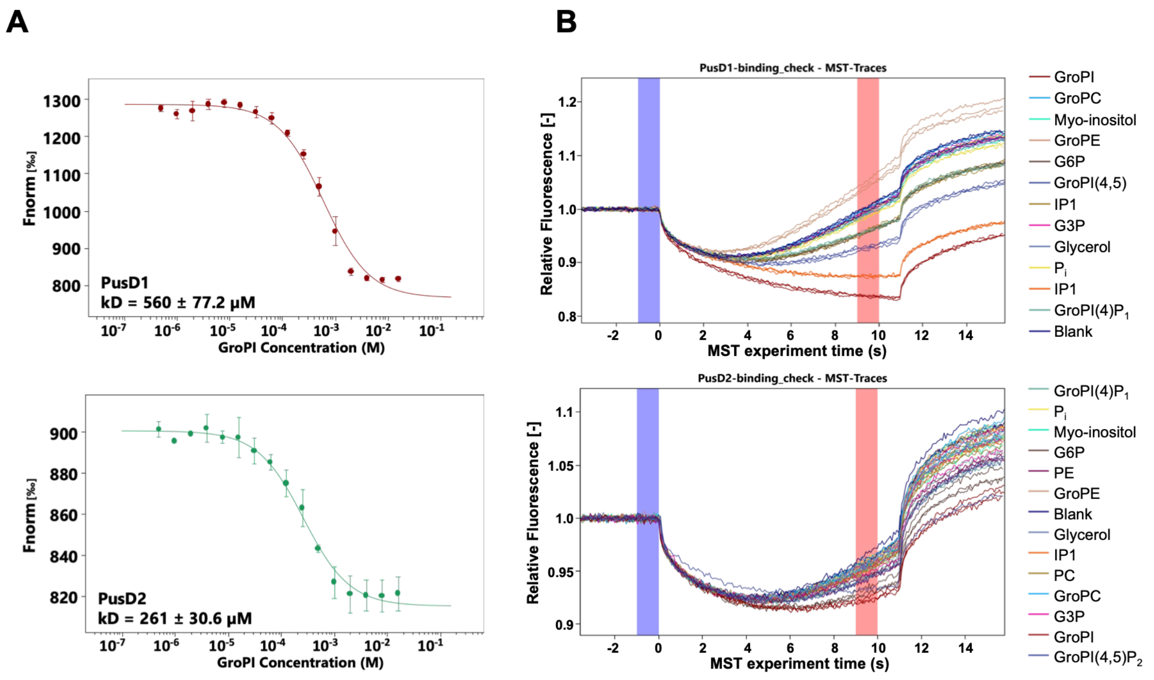


**Figure S13. Ligand-PusD binding interactions. A)** Microscale thermophoresis titration (max = 16 mM) for recombinant PusD1 (upper, red) and PusD2 (lower, green) interaction with GroPI. Dissociation constants ($K_d$)were estimated using the $K_d$ fit model within the Nanotemper MO.AffinityAnalysis software. Error bars represent standard error of the mean of quadruplicate measurements. **B)** Raw fluorescence traces for binding screens of each recombinant PusD protein against a diverse set of substrates (1 mM). Phosphorylated and non-phosphorylated substrates were screened. For both PusD proteins, no interaction with non-phosphorylated substrates, including myo-inositol and glycerol, or inorganic phosphate ($P_i$) was detected. For PusD1, the largest shifts observed were when incubated with inositol 1-phosphate (IP1) and glyceorphosphorylinositol (GroPI). Binding interactions between the mono- and bis-phosphorylated GroPI derivatives GroPI(4)$P_1$ and GroPI(4,5)$P_2$ as well as inositol triphosphate (IP3) was also detected. NB; IP3 was not tested for PusD2.

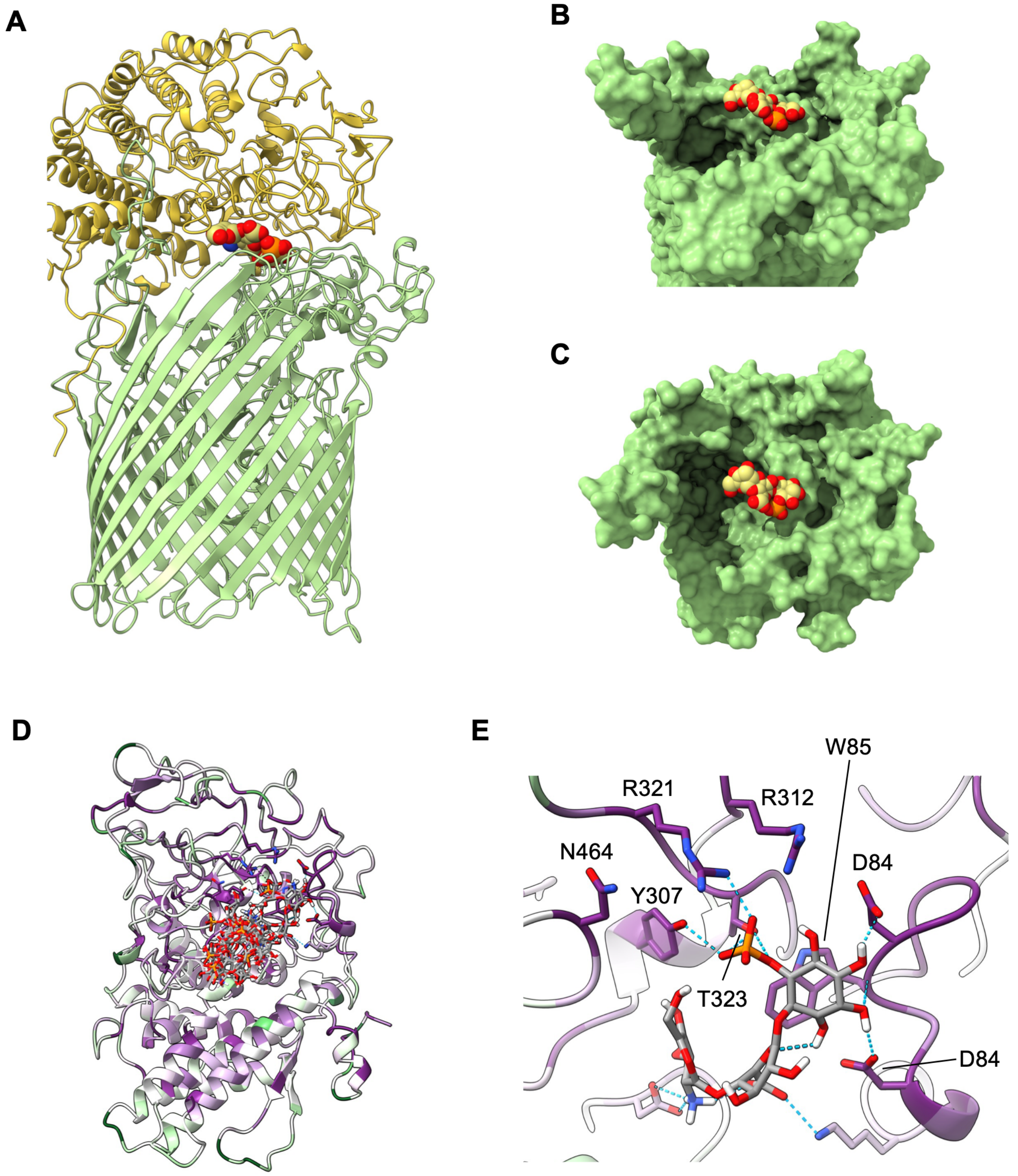


**Figure S14. Simulated binding of inositol phosphoglycan (IPG) in PusCD1. (A)** Atomic model of the closed PusCD1 complex with IPG, the potential metabolites of glycosyl inositolphosphoceramide (GIPC) post-hydrolysis by GIPC-PLC (Fjoh_4173) placed into the ligand binding cavity in relation to the IP1 moiety. The position of IPG was based on the experimental orientation of the inositol 1-phosphate moiety. **(B-C)** Surface topology of PusC1 illustrating the solvent-filled cavity. **D)** IPC docked with PusD1 using Autodock Vina (webina) showing the eight most likely poses[7]. **E)** The pose generated by mode 2 (affinity -6.132 kcal/mol) is shown, representing the most likely orientation of the IPG ligand based on our experimental data relating to IP1 binding. PusD1 residues are coloured based on ConSurf evolutionary conservation mapping[8], as per **Fig. 7**.

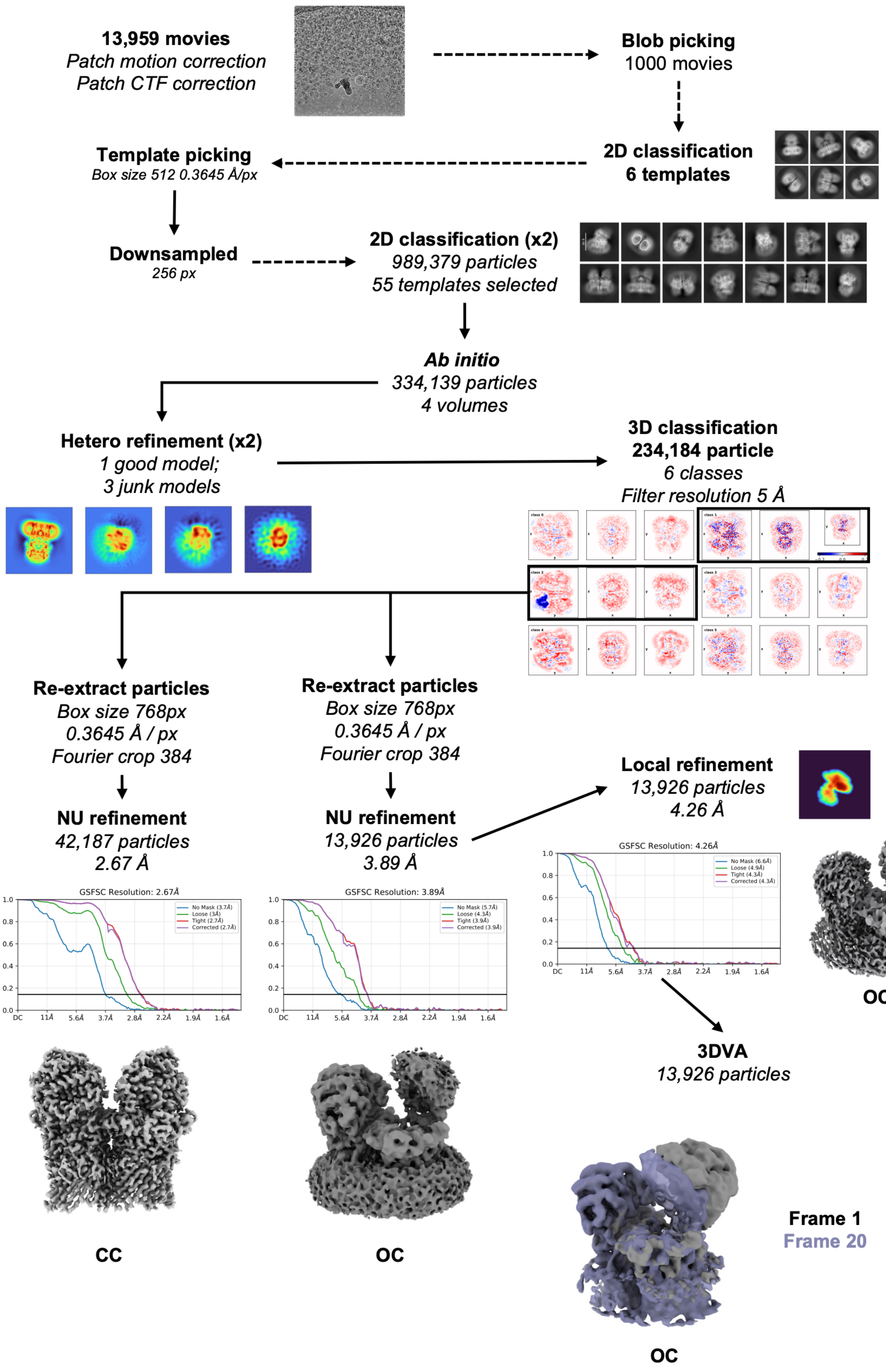


**Figure S15. Single-particle cryo-EM data processing workflow and resolution estimation of PusCD1.** CryoSPARC v4 was used for all steps. Mask refinement was performed in ChimeraX 10.1.1 The open-closed (OC) and closed-closed (CC) were refined from the same dataset. 3D variable analysis (3DVA) was used to investigate PusD1 lid mobility. Local refinement of the OC using a focused mask centred over the hinge and open PusD1 lid.

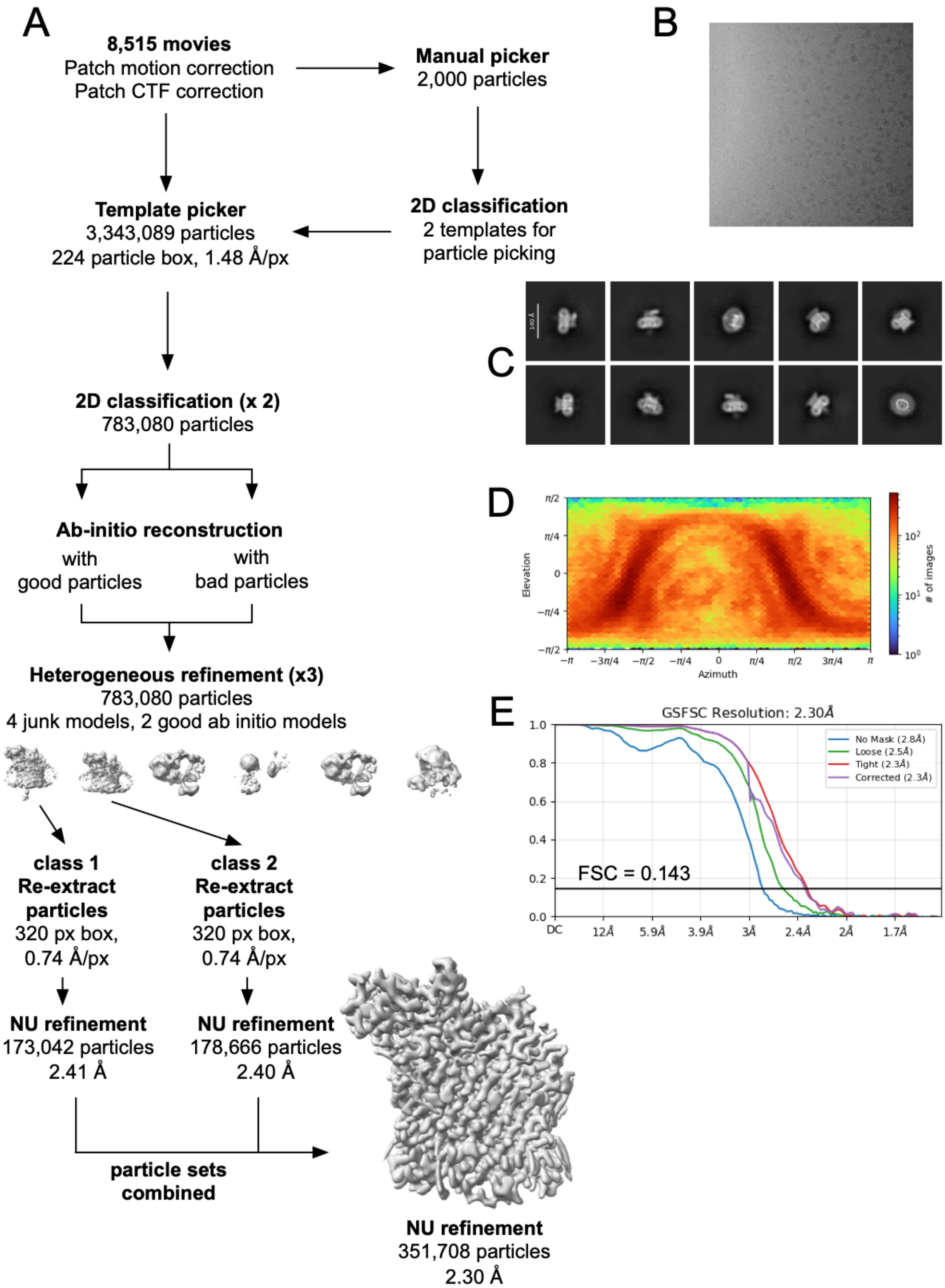


**Figure S16. Single-particle cryo-EM processing workflow and resolution estimation of the monomeric PusC1 A)** Schematic of the cryo-EM data processing pipeline using CryoSPARC v4. **B)** Example micrograph. **C)** 2D classes. **D)** Particle orientation distribution (azimuth vs. elevation) of the particles included in the final 3D reconstruction. **E) Fourier Shell Correlation (FSC) curves.** Gold-standard FSC (GSFSC) resolution estimation curves.
